# Evolutionary rescue of rare human genetic disorders in yeast models

**DOI:** 10.64898/2026.11.23.768747

**Authors:** Brooke Dubyna, Gregory I. Lang

## Abstract

Several thousand human disease-associated genes have clear yeast orthologs, making yeast an ideal model system for advancing a basic understanding of human disease. We used CRISPR-Cas9 editing to generate 51 yeast strains with 11 human disease-associated variants, focusing on human disease genes involved in protein glycosylation, ribosome maturation, mitochondrial function, and RNA regulation. We find that most of these alleles recapitulate as fitness defects in yeast. To enrich for suppressors, we evolved these populations for 1,000 generations and performed whole genome sequencing. We identified both known and previously unknown suppressor mutations. For example, we identify mutations in *DPM1* and *RPS9B* that compensate for disease phenotypes of *ALG7* and *SDO1*, respectively. Reconstructed strains harboring evolved suppressor mutations and the original disease alleles demonstrate that these suppressors rescue fitness and may compensate for defects in molecular function. By exerting continuous selective pressure over thousands of generations, experimental evolution allows us to select for suppressor mutations, including those with modest fitness effects or occurring at lower frequencies, which are underrepresented in traditional genetic screens. Our results map local genetic interaction networks centered around disease-associated alleles, providing greater insights into disease phenotypes and informing potential targeted therapies for disease management in patients with rare disorders.

## INTRODUCTION

Rare diseases collectively represent a substantial global health burden (Nguengang Wakap et al., 2020; Paltenghi-Peceva et al., 2026). Individuals affected by rare diseases often face significant challenges in accessing effective treatment options beyond symptomatic care. It is, therefore, critical to gain a better mechanistic understanding of the molecular bases of rare diseases. One strategy is to identify genetic suppressors—modifier genes interacting with disease-associated genes—which can serve as novel avenues for therapeutic interventions.

Model organisms offer a powerful tool for understanding the molecular basis of human disease, validating genetic interactions, and developing therapeutic strategies in systems with genetic tractability, large population sizes, and controlled experimental conditions (Hmeljak & Justice, 2019). With a fast doubling time, small genome size, and expansive availability of genomics tools, *Saccharomyces cerevisiae* has proven to be an effective model for studying eukaryotic cell biology (Botstein & Fink, 2011). Approximately 4,000 human genes have conserved orthologs in yeast, often performing the same cellular roles across organisms. This conservation enables the use of yeast strains containing human gene replacements or analogous mutations for functional, high-throughput studies of human biology and disease (Kachroo et al., 2022; Laurent et al., 2016).

Yeast experimental evolution can reveal genetic interactions by leveraging the power of natural selection to continuously enrich for beneficial mutations in replicate populations over hundreds or thousands of generations. While traditional suppressor screens tend to focus on the effects of gene deletions, we can evolve strains containing naturally occurring variants—including those in essential genes—such as nonsense and missense mutations, small insertions and deletions, or partial loss-of-function mutations. Further, this approach allows even modest fitness gains, such as those conferred by weak compensatory mutations, to be reproducibly detected. Combining experimental evolution with whole-genome sequencing allows us to identify and validate mutations that compensate for the deleterious effects of human disease-associated alleles (Vignogna et al., 2021, 2022).

Previously, we applied this approach to yeast strains containing human disease-associated alleles of *PMM2*, a gene associated with congenital disorder of glycosylation (CDG), type Ia, introduced into its yeast ortholog, *SEC53*. Using experimental evolution, we identified an overrepresentation of mutations in other CDG-associated genes: *PGM1*, *ALG7*, *ALG9*, and *ALG12*. Further characterization of evolved *PGM1* and *ALG9* mutations demonstrated allele-specific rescue of defects in glycosylation and growth. These findings highlight the strength of experimental evolution in identifying previously unknown genetic interactions and allele-specific suppressors of disease mutations (Vignogna et al., 2022).

Here, we applied experimental evolution to yeast strains with 11 additional human disease-associated allele-replacements, representing five rare genetic disorders. We evolved 377 individual populations, consisting of replicates of 51 mutant and three wildtype strains, for 1,000 generations and sequenced evolved clones to identify mutations that arose more often than would be expected to occur by chance. We found that mutations often occurred in genes functioning within the same pathways as their respective ancestral disease gene of interest. We demonstrate that evolved mutations of *DPM1* and *PSA1* compensate for growth defects and restore glycosylation deficiencies in *alg7* backgrounds. In *sdo1* backgrounds, we show allele-specific rescue of growth defects from mutations in *RIA1*, *RPS9B*, and *TIF6*. Together these results extend our previous work showing that experimental evolution can uncover allele-specific suppressors of human disease-associated variants.

## RESULTS

### Human disease-associated variants reduce growth rate in yeast

We selected six disease-associated human genes with 1:1 orthologs in yeast, each associated with rare, autosomal recessive genetic disorders. These genes are involved in a range of core, conserved biological processes such as protein glycosylation, ribosome maturation, mitochondrial function, and RNA modification. We chose common mutant alleles associated with each disease, where there is clinical evidence correlating the mutations to disease phenotypes in patients (Abdelrahman et al., 2018; Boocock et al., 2003; De Meirleir et al., 2004; Ng et al., 2018; Poyau et al., 2000; Shaheen et al., 2019; Teraoka et al., 1999). Across our six genes of interest, we selected 11 variants (Table 1) and constructed yeast strains by making the yeast-equivalent mutations through CRISPR-Cas9 editing. We generated a total of 51 human disease strains and three wildtype strains, including haploids of each mating type and both heterozygous and homozygous diploids. For genes for which we had multiple variants, we also constructed compound heterozygotes.

**Table 1.**

| Human Disease | Human Gene | Yeast Ortholog | Human Variant | Yeast Equivalent | Biological Process |
|---|---|---|---|---|---|
| Congenital disorder of glycosylation, type Ij | *DPAGT1* | *ALG7* | Y140C<br>R301C<br>R301H | Y171C<br>R336C<br>R336H | Protein N-linked glycosylation |
| Complex V deficiency, nuclear type 1 | *ATPAF2* | *ATP12* | W94R | W103R | Mito. ATP synthase complex assembly |
| Pseudouridine synthase deficiency | *PUS3*<br>*PUS7* | *DEG1*<br>*PUS7* | S394Cfs<br>D503Y | *deg1*Δ<br>D478Y | Pseudouridine synthesis |
| Shwachman-Diamond syndrome 1 | *SBDS* | *SDO1* | E44G<br>K62* | E44G<br>K62* | Cytosolic ribosome assembly |
| Leigh syndrome (Complex IV), nuclear type 1 | *SURF1* | *SHY1* | G124E<br>I246T<br>Y274D | G137E<br>F249T<br>Y344D | Mito. respiratory chain complex IV assembly |

While these disease-associated mutations tend to be highly pleiotropic in humans, manifesting in complex neurological or developmental phenotypes, equivalent disease alleles in yeast often result in slow growth (Kachroo et al., 2022; Laurent et al., 2020). To determine whether our variants of interest altered growth in yeast, we first performed tetrad dissections of heterozygous diploids (Fig. 1). In some cases, we observed a clear 2:2 segregation in colony sizes on YPD plates, representative of two slow-growing spores containing the disease allele and two wildtype spores displaying normal growth, while others had more subtle or no discernable visual differences in colony sizes. To quantitatively measure growth rate, we measured optical density of the spores in liquid cultures over 24 hours (Fig. 1). Most strains, including ones which previously showed little to no difference in colony sizes on plates, displayed 2:2 segregation patterns in their growth rates.

**Fig. 1.**
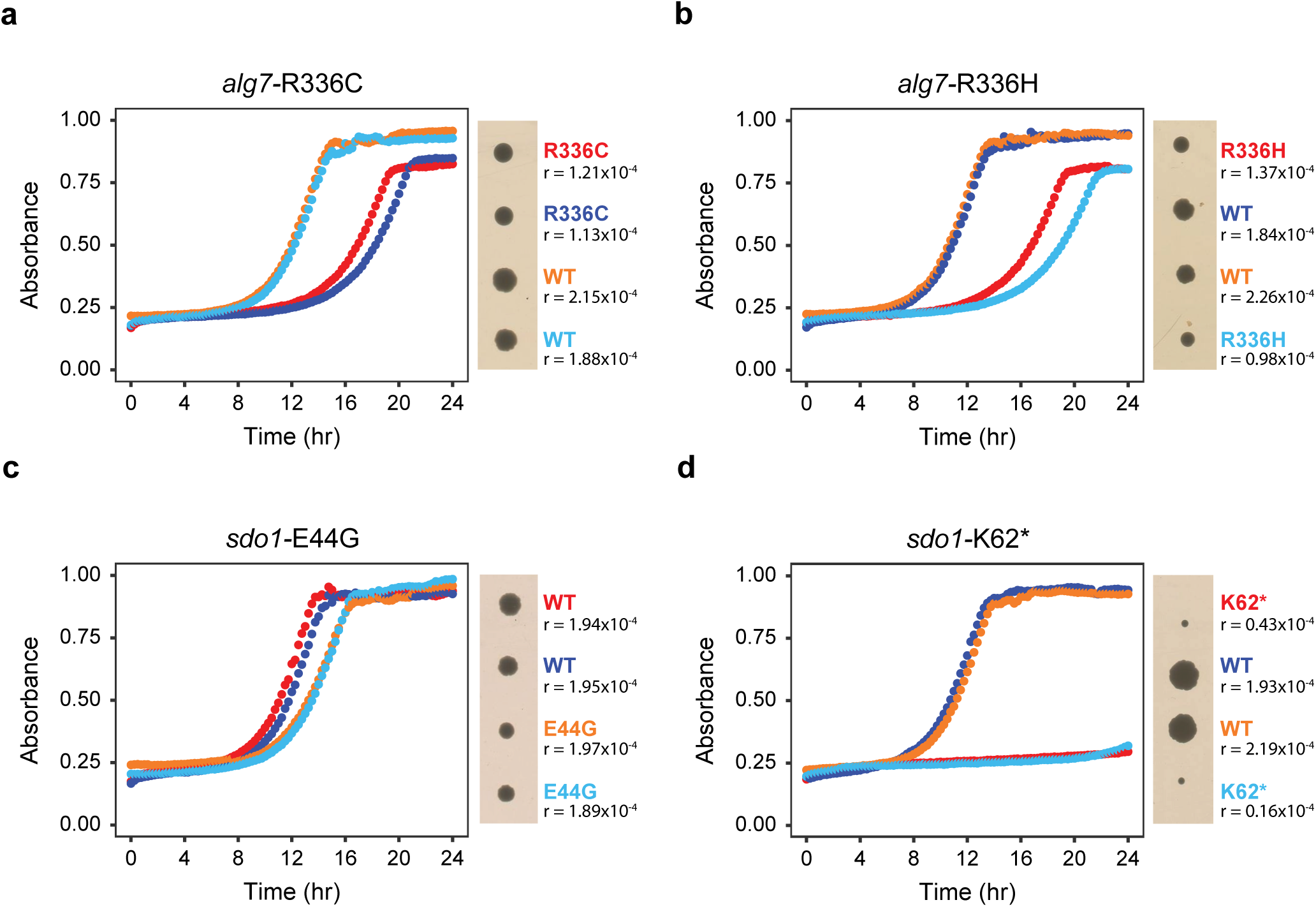
Growth curves measuring absorbance (OD_600_) over 24 hours for a subset of disease alleles: a) *alg7*-R336C, b) *alg7*-R336H, c) *sdo1*-E44G, and d) *sdo1*-K62*. The four colored lines each represent the growth of an individual spore of a four-spore tetrad from a heterozygous diploid strain. Growth rate (r) is indicated for each curve. Differences in growth between the four spores are also observed by colony growth on YPD plates (to the right of each graph).This subset all show distinct 2:2 segregation of wildtype and disease-associated alleles, with spores inheriting disease-associated alleles displaying slower growth compared to wildtype spores.

To more precisely quantify fitness effects of each disease allele in yeast, we performed a competitive fitness assay, competing each strain against a fluorescently labeled wildtype ancestor (Table 2). If a human disease allele performs a conserved role in yeast and humans, we expect to observe fitness defects in the yeast harboring this variant. We found that the majority of haploids and homozygous diploids with disease allele replacements showed decreased fitness relative to wildtype populations. For instance, *sdo1*-K62* was highly deleterious, resulting in a fitness effect of -17.82 ± 2.28% (mean ± SD) in homozygous diploids. However, some disease alleles, including *alg7*-Y171C, *atp12*-W103R, *shy1*-F249T, and *shy1*-Y344D, did not cause fitness defects in yeast and instead performed comparably to wildtype populations (1.42 ± 0.10%, 0.26 ± 0.12%, 0.40 ± 0.17%, and -0.03 ± 0.15%, respectively), despite strong clinical evidence of a disease-associated phenotype in humans.

**Table 2.**
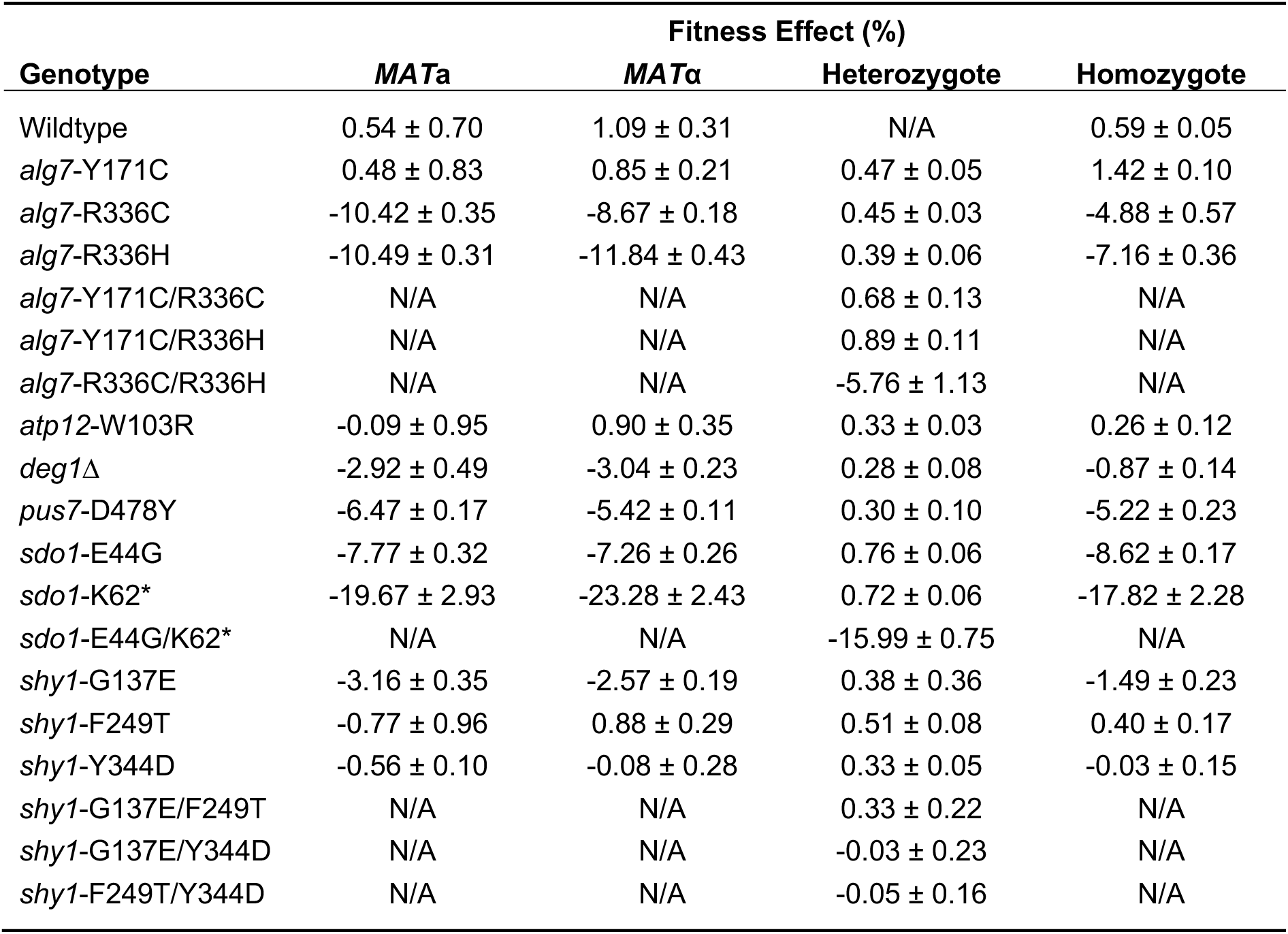

|  | Fitness Effect (%) |  |  |  |
|---|---|---|---|---|
| **Genotype** | ***MAT*a** | ***MAT*α** | **Heterozygote** | **Homozygote** |
| Wildtype | 0.54 ± 0.70 | 1.09 ± 0.31 | N/A | 0.59 ± 0.05 |
| *alg7*-Y171C | 0.48 ± 0.83 | 0.85 ± 0.21 | 0.47 ± 0.05 | 1.42 ± 0.10 |
| *alg7*-R336C | -10.42 ± 0.35 | -8.67 ± 0.18 | 0.45 ± 0.03 | -4.88 ± 0.57 |
| *alg7*-R336H | -10.49 ± 0.31 | -11.84 ± 0.43 | 0.39 ± 0.06 | -7.16 ± 0.36 |
| *alg7*-Y171C/R336C | N/A | N/A | 0.68 ± 0.13 | N/A |
| *alg7*-Y171C/R336H | N/A | N/A | 0.89 ± 0.11 | N/A |
| *alg7*-R336C/R336H | N/A | N/A | -5.76 ± 1.13 | N/A |
| *atp12*-W103R | -0.09 ± 0.95 | 0.90 ± 0.35 | 0.33 ± 0.03 | 0.26 ± 0.12 |
| *deg1*∆ | -2.92 ± 0.49 | -3.04 ± 0.23 | 0.28 ± 0.08 | -0.87 ± 0.14 |
| *pus7*-D478Y | -6.47 ± 0.17 | -5.42 ± 0.11 | 0.30 ± 0.10 | -5.22 ± 0.23 |
| *sdo1*-E44G | -7.77 ± 0.32 | -7.26 ± 0.26 | 0.76 ± 0.06 | -8.62 ± 0.17 |
| *sdo1*-K62\* | -19.67 ± 2.93 | -23.28 ± 2.43 | 0.72 ± 0.06 | -17.82 ± 2.28 |
| *sdo1*-E44G/K62\* | N/A | N/A | -15.99 ± 0.75 | N/A |
| *shy1*-G137E | -3.16 ± 0.35 | -2.57 ± 0.19 | 0.38 ± 0.36 | -1.49 ± 0.23 |
| *shy1*-F249T | -0.77 ± 0.96 | 0.88 ± 0.29 | 0.51 ± 0.08 | 0.40 ± 0.17 |
| *shy1*-Y344D | -0.56 ± 0.10 | -0.08 ± 0.28 | 0.33 ± 0.05 | -0.03 ± 0.15 |
| *shy1*-G137E/F249T | N/A | N/A | 0.33 ± 0.22 | N/A |
| *shy1*-G137E/Y344D | N/A | N/A | -0.03 ± 0.23 | N/A |
| *shy1*-F249T/Y344D | N/A | N/A | -0.05 ± 0.16 | N/A |

Additionally, we found that diploids often performed slightly better than their haploid counterparts, indicating that disease alleles had slightly greater fitness costs in haploids (Table 2). For example, *MAT***a** *alg7*-R336H populations displayed a fitness effect of -10.49 ± 0.31%, compared to -7.16 ± 0.36% for homozygous diploids. As expected for recessive inheritance, all heterozygous diploids had no fitness defects since they possess one wildtype allele. Compound heterozygotes displayed fitness effects intermediate of those observed for each of the individual alleles. For example, *sdo1*-E44G and *sdo1*-K62* had fitness effects of -8.62 ± 0.17% and -17.82 ± 2.28%, respectively. The compound heterozygote had a fitness effect of -15.99 ± 0.75%, not statistically different from the additive expectation of -13.22 ± 2.29% (p = 0.068, two-sided Welch-type t-test). In cases where a variant had no effect on fitness, compound heterozygotes containing that variant also displayed no defects.

### Laboratory evolution restores fitness of strains carrying disease-associated alleles

We evolved replicates of each strain for 1,000 generations in nutrient-rich media, performing competitive fitness assays at generations 300, 700, and 1,000 (Fig. 2). Compared to starting fitness effects at generation 0, nearly all populations increased in fitness by generation 300, often achieving near-wildtype fitness levels, and continuing to gradually increase in fitness over subsequent generations. These findings suggest that the disease strains acquired compensatory mutations throughout the evolution experiment which served to rescue fitness defects.

**Fig. 2.**
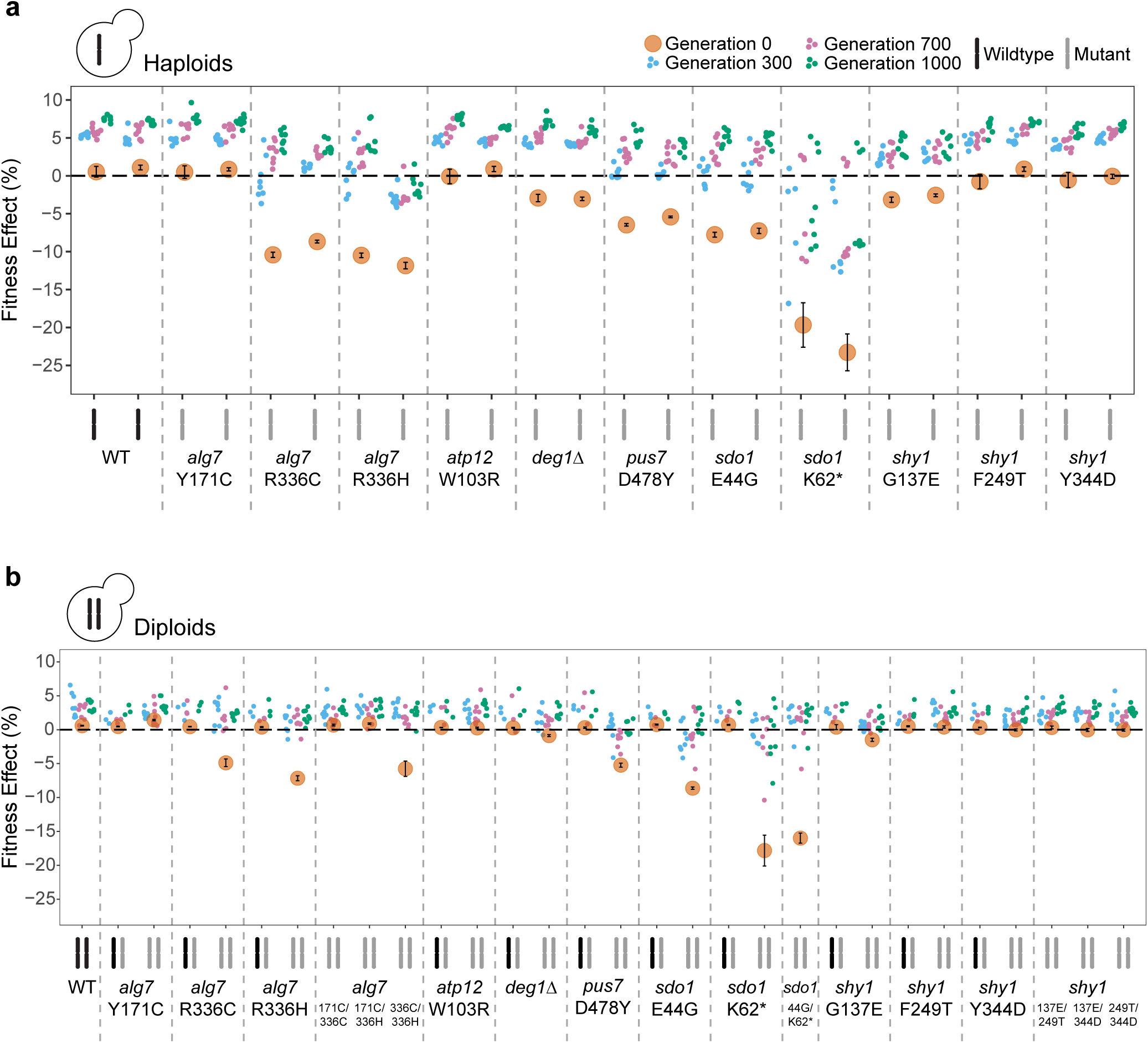
Fitness effects of each population throughout the evolution experiment measured by the competitive fitness assay. Since the ancestral populations for each variant are replicates, Generation 0 populations are averaged into one data point (orange) with standard deviations shown. Points for generations 300, 700, and 1,000 represent individually evolving populations. A fitness effect at 0% along the horizontal dashed line is indicative of wildtype fitness; fitness effects above or below represent a fitness advantage or disadvantage, respectively, compared to wildtype. a) Haploid strains include both *MAT***a** and *MAT*α, shown in that order. Haploid *sdo1*-K62* fitness effects for Generation 0 were measured during later reconstructions using haploid spores inheriting the K62* allele, as the original strains at Generation 0 were too rapidly outcompeted to accurately measure fitness effects. b) Diploid strains include heterozygotes, homozygotes, and compound heterozygotes

To identify potential compensatory mutations, we isolated single clones from each evolved population at generation 1,000 and performed whole genome sequencing. Putative adaptive targets of selection are genes with more mutations than expected to occur by random chance (Fig. 3). Across all 377 evolved populations, we identified 1336 mutations, 822 of which were nonsynonymous mutations within coding regions (Supplemental Dataset 2). We identified mutations falling into two categories: common targets of selection and mutations unique to disease-associated genetic backgrounds. Common targets include *IRA1*, *IRA2*, and *WHI2*, in which mutations were not specific to the disease genes of interest. These genes accrued mutations across several different experimental groups and are frequently observed in other laboratory evolution experiments (Fisher et al., 2018; Johnson et al., 2021; Lang et al., 2013; Vignogna et al., 2022). Mutations in these common targets confer a general fitness advantage and likely do not compensate for disease phenotypes.

**Fig. 3.**
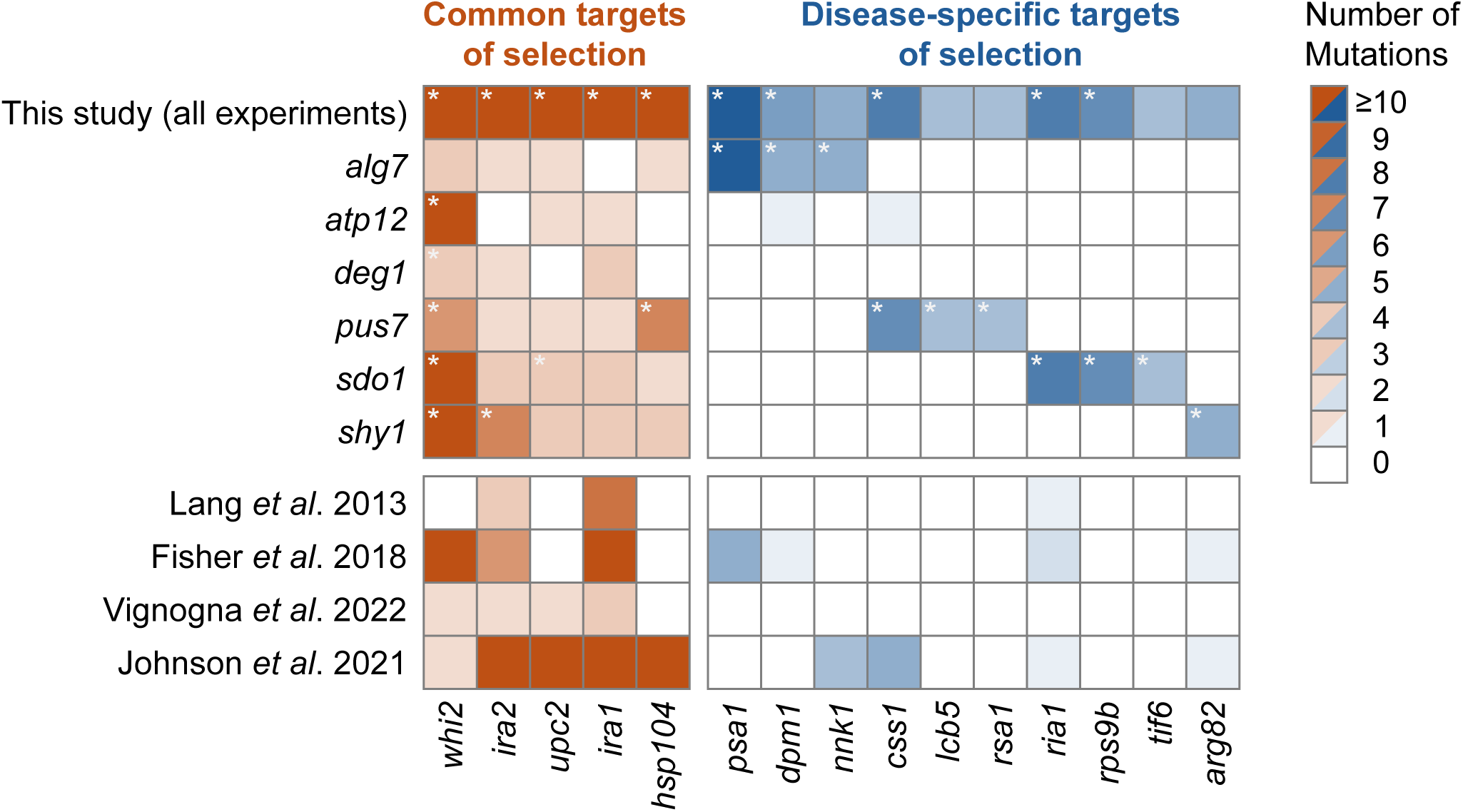
Heatmap showing the number of nonsynonymous mutations per gene that arose during the evolution experiment. Genes with a significant number of mutations are indicated by asterisks (*). Genes are separated by common targets and disease-specific targets of selection. For comparison, we include data from previously reported evolution experiments using similar experimental conditions with different strain backgrounds.

### Putative allele-specific suppressors are identified by overrepresentation

Genes that are mutated more often than expected by chance in a disease-associated background are potentially compensating for that disease allele. In the seven *alg7* backgrounds which demonstrated fitness defects—*alg7*-R336C and *alg7*-R336H haploids, homozygotes, and their compound heterozygote—we found more mutations than expected by chance in *PSA1* (11 nonsynonymous mutations, Poisson probability = 7.0 x 10^-31^), *DPM1* (5, P = 1.0 x 10^-13^), and *NNK1* (5, P = 5.1 x 10^-11^). Of these, there were eight distinct mutations in *PSA1*, two in *DPM1*, and one in *NNK1*. One *alg7*-R336H population acquired two different mutations in *PSA1*. In some cases, as with *NNK1*, the same mutation was observed across multiple populations, suggesting it may have been present in the ancestral population and subsequently rose to high frequency across populations. In contrast, instances of multiple distinct mutations are indicative of mutations arising independently across populations. Further, *PSA1* and *DPM1* gene products function in the same cellular pathway as *ALG7*, protein glycosylation.

Similarly, *sdo1* mutants displayed an enrichment of mutations in genes involved in the same ribosomal maturation pathway as *SDO1*: *RPS9B* (7 nonsynonymous mutations, Poisson probability = 6.3 x 10^-17^), *RIA1* (8, P = 1.9 x 10^-15^), and *TIF6* (4, P = 8.9 x 10^-10^). There were two distinct mutations in *RPS9B*, seven in *RIA1*, and four in *TIF6*. Of note, mutations in *RPS9B* arose only in *sdo1*-K62* backgrounds, while *RIA1* mutations were exclusive to *sdo1*-E44G populations, suggesting that putative suppressors may be allele-specific.

Additionally, all observed mutations in these genes found in *alg7* and *sdo1* backgrounds were missense mutations, with the exception of one conservative in-frame deletion in *RPS9B*. Given we have between one and eight independent mutations in each suppressor gene, we used a Bayesian approach to assess whether these mutations were consistent with loss-of-function (LOF) or non-loss-of-function (non-LOF) effects (Martínez & Lang, 2023). In each case, they supported the hypothesis that selection favored non-LOF mutations, with posterior probabilities of non-LOF ranging from 0.56 to 0.996 (log_10-_likelihood ratio (LLR) = -0.32 to -2.57). This finding is consistent with these mutations occurring primarily in essential genes in which LOF mutations may not be tolerated.

In *pus7* mutants, we observed mutations in *CSS1* (6 nonsynonymous mutations, Poisson probability = 6.1 x 10^-13^), *RSA1* (4, P = 5.2 x 10^-10^), and *LCB5* (4, P = 5.5 x 10^-9^). As noted previously with *NNK1* in *alg7* backgrounds, each of these genes had one common mutation across multiple populations, rather than multiple distinct mutations, suggesting that these mutations were present at low frequencies in the ancestral cultures and subsequently fixed in several populations during the evolution experiment. Unlike putative suppressors of *alg7* and *sdo1*, none of these genes function in similar pathways.

### Genetic reconstructions validate putative suppressor mutations

To validate putative suppressors of *alg7* and *sdo1* for their ability to compensate for disease phenotypes, we reconstructed diploid strains that were heterozygous for an original disease-allele and a corresponding evolved mutation using CRISPR-Cas9 allele replacement. For each putative suppressor gene, we selected two evolved mutations for reconstructions, with the exception of *NNK1*, in which only one variant arose multiple times across the evolution experiment. However, we also included a full gene deletion of *NNK1*, as this was one of two non-essential putative suppressor genes, in order to determine whether complete loss of function was sufficient to compensate for defects. Despite multiple attempts, we were unable to successfully generate the other non-essential gene deletion of *RPS9B*, as well as one variant in the background of *sdo1*-E44G, *tif6*-F7L.

We sporulated and dissected tetrads from each of these heterozygote reconstructions. If suppression occurred, haploid spores inheriting both the disease and suppressor alleles would form larger colonies with normal or intermediate growth, indicating that the suppressor was rescuing growth defects, while those inheriting the disease allele alone would continue showing inhibited growth. We expect three types of segregation patterns of large to small colonies: 2:2, 3:1, and 4:0. Assuming no genetic linkage, these three segregation patterns are expected to follow a 1:4:1 ratio. In instances of no suppression, we would observe only the 2:2 segregation of the disease allele, as previously noted without a suppressor present. Based on these ratios of colony growth, we found that *alg7*-R336C and *alg7*-R336H were suppressed by each variant of *DPM1* and *PSA1* (Fig. 4, Supplemental Dataset 3). For instance, *alg7*-R336C/*dpm1*-A185D tetrads displayed segregation patterns consistent with suppression (LLR = 26.85, exact multinomial p-value = 6.3 x 10^-29^). While a full deletion of *NNK1* did not act as a suppressor (for *alg7*-R336C and *alg7*-R336H, respectively; LLR = -7.20, -0.91), *nnk1*-R326K demonstrated possible weak suppression with intermediate colonies appearing only slightly larger than those with the disease mutation alone (LLR = 5.74, 4.22; p = 2.4 x 10^-9^, 1.6 x 10^-7^).

**Fig. 4.**
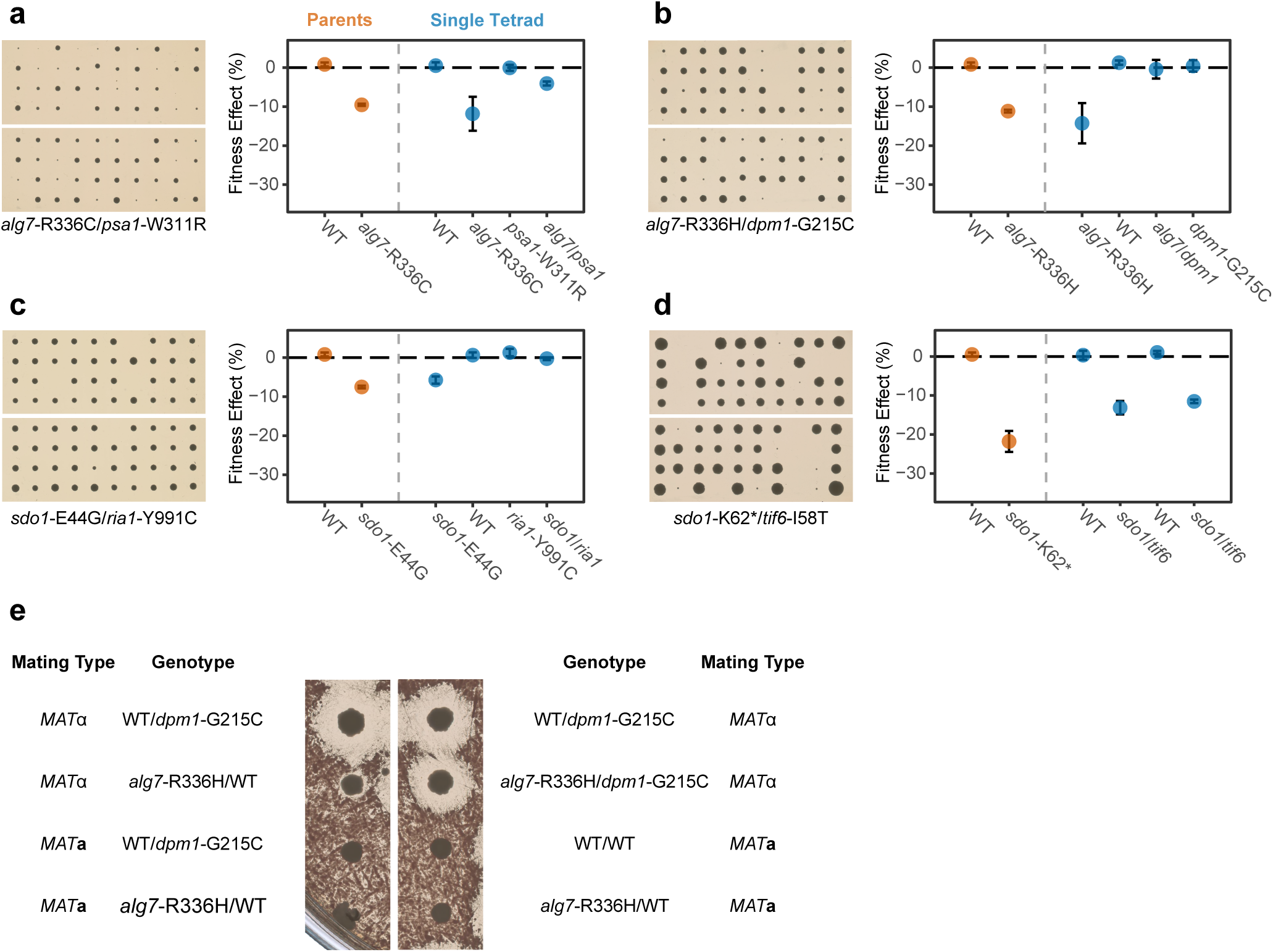
Tetrad dissections and fitness effects for a subset of heterozygous reconstructions containing ancestral disease-associated alleles and putative suppressors: a) *alg7*-R336C/*psa1*-W311R, b) *alg7*-R336H/*dpm1*-G215C, c) *sdo1*-E44G/*ria1*-Y991C, d) *sdo1*-K62*/- *tif6*-I58T. Plate pictures for tetrad dissections were taken after two days of growth, with the exception of d) which required six days for *sdo1*-K62* spores to become visible. Segregation patterns of large to small colonies were consistent with suppression. Fitness effects were measured by competitive fitness assay. We show four spores from one tetrad per cross, with error bars representing the standard error of slope. We include a comparison of wildtype and averaged *MAT***a** and *MAT*α haploids from Generation 0 of the evolution experiment, with error bars indicated the standard deviation of the mean. Spores inheriting both the ancestral disease-associated allele and the putative suppressor allele show increased fitness relative to those with the disease-associated allele alone. e) Halo assay with *alg7*-R336H/*dpm1*-G215C tetrads. *MAT*α spores create a halo on a lawn of *MAT***a** cells. Spores inheriting *alg7*-R336H display a decreased halo size, while those which also inherit the suppressor allele (*alg7*-R336H/*dpm1*-G215C) have halos restored to near-wild-type halo sizes.

In *sdo1*-K62* strains, each variant of *RPS9B* and *TIF6*, but neither of the *RIA1* mutations, served as a suppressor. Reconstructions of *sdo1*-K62*/*rps9b*-K92Δ, for example, supported suppression (LLR = 24.91, p = 7.4 x 10^-27^), while *sdo1*-K62*/*ria1*-P368S tetrads lacked suppression (LLR = -5.16). Since *sdo1*-E44G had less obvious differences in colony sizes when comparing wildtype and mutant growth, we were unable to differentiate segregation patterns from growth on plates.

### Extent of suppression varies by disease-associated allele and suppressor

To identify the extent of suppression, we measured the fitness of each spore using the competitive fitness assay to determine whether the presence of a putative suppressor mutation increased fitness in the background of disease mutations (Fig. 4, Supplemental Fig. 1). We again found that variants of *DPM1* and *PSA1* were compensatory in both *alg7*-R336 backgrounds. For instance, in *alg7*-R336C/*dpm1*-A185D reconstructions, spores inheriting both mutations exhibited a mean fitness effect of -1.24 ± 1.16% (mean ± SD, n = 4), compared to -13.47 ± 1.81% (n = 4) for *alg7*-R336C/*DPM1* and 0.62 ± 0.50% (n = 4) for wildtype (or *ALG7/dpm1*-A185D), indicating that the suppressor conferred nearly full rescue to wildtype fitness levels with approximately 87% suppression (Table 3, Supplemental Fig. 1). Similarly, in *alg7*-R336H/*psa1*-W311R reconstructions, we observed a mean fitness effect of -6.10 ± 1.62% (n = 5) for spores with both mutations, compared to -16.26 ± 1.31% (n = 3) for *alg7*-R336H/*PSA1*. In this case, *psa1*-W311R conferred partial rescue of the disease allele, restoring approximately 60% of the fitness defect. In contrast, *nnk1*-R326K and *nnk1*Δ showed no clear evidence of suppression, with fitness following 2:2 segregation patterns of wildtype and mutant fitness levels.

**Table 3.**
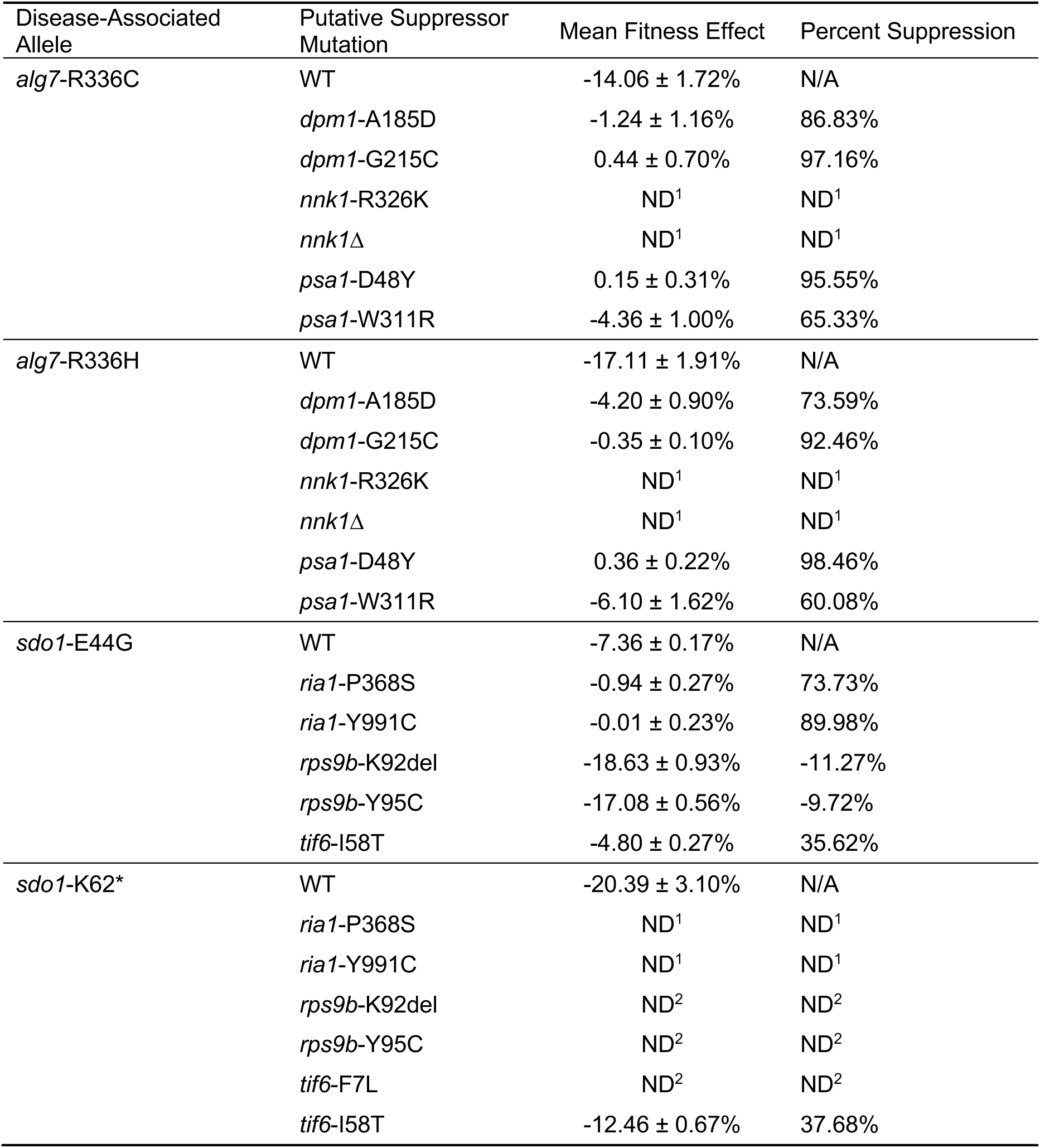
^1^ Spores inheriting the disease-associated allele alone or in combination with the putative suppressor had fitness effects indistinguishable from one another. ^2^ Fitness effects of spores inheriting *sdo1*-K62* in combination with a weak suppressor were indistinguishable from those inheriting the K62* allele alone. These spores had severe fitness defects, causing them to be outcompeted to the point of extinction within five to ten generations of beginning the competitive fitness assay. Differences in fitness were apparent on YPD plates, where double mutant spores formed larger colonies than those inheriting the disease-associated allele alone.

| Disease-Associated Allele | Putative Suppressor Mutation | Mean Fitness Effect | Percent Suppression |
|---|---|---|---|
| *alg7*-R336C | WT | -14.06 ± 1.72% | N/A |
|  | *dpm1*-A185D | -1.24 ± 1.16% | 86.83% |
|  | *dpm1*-G215C | 0.44 ± 0.70% | 97.16% |
|  | *nnk1*-R326K | ND<sup>1</sup> | ND<sup>1</sup> |
|  | *nnk1*Δ | ND<sup>1</sup> | ND<sup>1</sup> |
|  | *psa1*-D48Y | 0.15 ± 0.31% | 95.55% |
|  | *psa1*-W311R | -4.36 ± 1.00% | 65.33% |
| *alg7*-R336H | WT | -17.11 ± 1.91% | N/A |
|  | *dpm1*-A185D | -4.20 ± 0.90% | 73.59% |
|  | *dpm1*-G215C | -0.35 ± 0.10% | 92.46% |
|  | *nnk1*-R326K | ND<sup>1</sup> | ND<sup>1</sup> |
|  | *nnk1*Δ | ND<sup>1</sup> | ND<sup>1</sup> |
|  | *psa1*-D48Y | 0.36 ± 0.22% | 98.46% |
|  | *psa1*-W311R | -6.10 ± 1.62% | 60.08% |
| *sdo1*-E44G | WT | -7.36 ± 0.17% | N/A |
|  | *ria1*-P368S | -0.94 ± 0.27% | 73.73% |
|  | *ria1*-Y991C | -0.01 ± 0.23% | 89.98% |
|  | *rps9b*-K92del | -18.63 ± 0.93% | -11.27% |
|  | *rps9b*-Y95C | -17.08 ± 0.56% | -9.72% |
|  | *tif6*-I58T | -4.80 ± 0.27% | 35.62% |
| *sdo1*-K62\* | WT | -20.39 ± 3.10% | N/A |
|  | *ria1*-P368S | ND<sup>1</sup> | ND<sup>1</sup> |
|  | *ria1*-Y991C | ND<sup>1</sup> | ND<sup>1</sup> |
|  | *rps9b*-K92del | ND<sup>2</sup> | ND<sup>2</sup> |
|  | *rps9b*-Y95C | ND<sup>2</sup> | ND<sup>2</sup> |
|  | *tif6*-F7L | ND<sup>2</sup> | ND<sup>2</sup> |
|  | *tif6*-I58T | -12.46 ± 0.67% | 37.68% |

Variants of *RIA1* were compensatory in *sdo1*-E44G, but not in *sdo1*-K62* backgrounds. For example, in *sdo1*-E44G/*ria1*-Y991C tetrads, double mutant spores displayed a mean fitness effect of -0.01 ± 0.23% (n = 3), while *sdo1*-E44G/*RIA1* spores scored -6.75 ± 0.74% (n = 5). This suppression conferred nearly full rescue, with approximately 90% fitness restoration. However, *sdo1*-K62*/*ria1*-Y991C reconstructions displayed no discernable differences between *sdo1*-K62*/*RIA1* and double mutants, with tetrads displaying the same 2:2 segregation patterns observed in the absence of a suppressor.

Conversely, *RPS9B* variants conferred improved growth in the *sdo1*-K62* background but displayed decreased fitness in wildtype (−13.36 ± 0.61%, n = 8; -9.36 ± 0.42%, n = 8; for *rps9b*-K92Δ and *rps9b*-Y95C, respectively) and *sdo1*-E44G backgrounds (−18.62 ± 0.93%, n = 4; -17.08 ± 0.56%, n = 4). Despite observing clear increases in colony sizes for *sdo1*-K62*/*rps9b*-K92Δ and *sdo1*-K62*/*rps9b*-Y95C double mutants compared to *sdo1*-K62*/*RPS9B*, indicating partial rescue of growth defects, we were unable to recapitulate this improved fitness in our competitive fitness assay.

*TIF6* variants, on the other hand, demonstrated compensatory effects in both *sdo1* backgrounds. Spores from reconstructions with *tif6*-I58T displayed mean fitness effects of -7.80 ± 0.75% (n = 3) for *sdo1*-E44G/*TIF6* and -4.80 ± 0.27% (n = 5) for *sdo1*-E44G/*tif6*-I58T, indicating a 36% fitness restoration. Similarly, *sdo1*-K62*/*tif6*-I58T yielded a fitness of -12.46 ± 0.67% (n = 6). None of the spores from this cross selected for fitness assays had a genotype of *sdo1*-K62*/*TIF6*, but in comparison to other *sdo1*-K62* spores with a mean fitness effect of -20.39 ± 3.10% (n = 10), *tif6*-I58T restored fitness by approximately 38%. Interestingly, this same variant appears slightly deleterious in wildtype backgrounds, with a fitness of -2.99 ± 0.28% (n = 11); however, *tif6*-F7L was indistinguishable from wildtype.

### Suppressors of *alg7* rescue molecular defects in protein glycosylation

Suppressors of disease-associated alleles could be acting to restore the molecular defect or compensate for it. For the *ALG7* disease alleles, we can distinguish between these possibilities by assaying α-factor secretion since prepro-α-factor glycosylation is necessary for efficient transport through the secretory pathway (Caplan et al., 1991). Consistent with this process, we observed that *alg7*-R336 mutants form small halos on a lawn of *MAT***a** cells (Fig. 4e, Supplemental Dataset 3). We found that double mutants with both the disease allele and *DPM1* or *PSA1* suppressors—*dpm1*-A185D, *dpm1*-G215C, *psa1*-D48Y, or *psa1*-W311R—often had halos restored to intermediate or near wildtype sizes, suggesting that the suppressors may be serving to resolve molecular defects in glycosylation.

## DISCUSSION

We generated yeast models of rare human diseases by selecting six disease-associated human genes with 1:1 orthologs in yeast, where molecular function is likely to be highly conserved (Gabaldón & Koonin, 2013). We chose known pathogenic variants in which the wildtype amino acid residue is conserved between humans and yeast. Nevertheless, four of the eleven clinically pathogenic variants fail to show a fitness defect in yeast. These differences can potentially be explained by the molecular mechanisms of the mutations and the conditions under which fitness was assessed. For example, a yeast-specific co-chaperone Fmc1 helps to maintain Atp12 solubility (Meulemans et al., 2010), and may therefore mask the effect of the *atp12*-W103R mutation. Both *ATP12* and *SHY1* are involved in mitochondrial function, and both *atp12*-W103R and *shy1*-Y344D have previously demonstrated growth defects on nonfermentable carbon sources, such as glycerol, but displayed normal growth on YPD (Bestwick et al., 2010; Meulemans et al., 2010). In our assay, however, *shy1*-F249T showed no defects even on nonfermentable carbon sources.

Within *ALG7*, two of the three alleles recapitulate a disease phenotype in yeast. Variants at residue R336 (R336C and R336H) phenocopied human disease by exhibiting large fitness defects, while *alg7*-Y171C appeared to have no measurable effect on growth rate. This difference between variants may be accounted for by the amino acid positioning and role in protein function. In the orthologous human protein, the equivalent R336 residue, R301, lies within the highly conserved active site, serving to stabilize the protein substrate, while Y140 (Y171 in yeast) is within a transmembrane domain (Dong et al., 2018; Ng et al., 2018). Mutations affecting enzymatic activity may, therefore, be more likely to recapitulate a disease phenotype than are mutations affecting protein stability.

Following 1,000 generations of laboratory evolution, all strains improved in fitness, with most being restored to wildtype fitness levels by generation 300, consistent with the expected timing of the first selective sweep (Lang et al., 2013). We continued the evolution experiment through 1,000 generations to allow mutations with more subtle fitness effects to arise, recognizing that early mutations may be common drivers of adaptation rather than background-specific suppressors (Fisher et al., 2018; Lang et al., 2013; Vignogna et al., 2022).

We identified *DPM1* and *PSA1* as suppressors of *alg7*, and *RIA1*, *RPS9B*, and *TIF6* as suppressors of *sdo1*. We were unable to validate *NNK1* as a suppressor of *alg7* despite being identified as a statistically-significant target of selection in the evolution experiment. Evolution can conceivably act on selection coefficients several orders of magnitude smaller than we can detect in the laboratory; therefore, *NNK1* may be a weak suppressor not picked up in our competitive fitness assay. We observe a rescue of α-factor secretion with *dpm1* and *psa1* suppressors in the *alg7* background, indicating that these mutations restore wildtype levels of protein glycosylation, something that we previously observed for *pgm1* suppressors in the *sec53* background (Vignogna et al., 2022).

We identified *RPS9B* and *TIF6* as suppressors of *sdo1*-K62* based on their ability to increase colony size in the *sdo1*-K62* background. However, with the exception of *tif6*-I58T, we did not observe measurable fitness gains in the competitive fitness assay. In these cases, the combination of the severe fitness defect of the *sdo1*-K62* mutation and the modest effects of the suppressor mutations resulted in both the original mutants and the suppressed strains being rapidly outcompeted by the reference strain, obscuring small fitness improvements. Compensation from weak suppressors in the background of severely deleterious mutations may, therefore, be better evaluated on plates rather than with a competitive fitness assay.

We find that genetic suppressors often fall in the same biological pathway as the disease-associated gene. *ALG7* and its suppressors *DPM1* and *PSA1* are all essential genes in the N-linked glycosylation pathway. The human orthologs of *PSA1* and *DPM1* (*GMPPB* and *DPM1*, respectively) are associated with various forms of muscular dystrophy-dystroglycanopathy and congenital disorder of glycosylation, type Ie, respectively (OMIM #615350-615352, 608799). *SDO1* and its suppressors *RIA1*, *RPS9B*, and *TIF6* function in ribosome maturation, with *RIA1* and *TIF6* each being essential genes. The human ortholog of *RIA1* (*EFL1*) is implicated in Shwachman-Diamond syndrome (OMIM #617941). This enrichment of mutations within the same pathway as the original disease-associated gene is in line with previous findings, in which *sec53* backgrounds accumulated mutations in other CDG homologs (Vignogna et al., 2022). In fact, approximately 36% of known suppression interactions reported in patients, cultured human cells, or animal models occur between genes of the same biological process (Paltenghi-Peceva et al., 2026).

Neither *DPM1* nor *PSA1* are known genetic or physical interactors of *ALG7* in yeast; however, *DPM1* was identified as an interactor of *ALG7* in *Drosophila melanogaster*, with inhibition of Dpm1 rescuing disease phenotypes (Dalton et al., 2022). We hypothesize that mutations altering function of Dpm1 or Psa1 rescue *alg7* phenotypes by altering flux of dolichol-phosphate, a substrate of both Dpm1 and Alg7, either increasing substrate availability for Alg7 or promoting glycosylation through the branch of the pathway involving Dpm1 and Psa1. Of note, all evolved mutations in *PSA1* occurred in residues within either the N- or C-terminal, which are the two major functional domains of the Psa1 protein in which mutations are typically associated with disease (Chompoopong & Milone, 2023). This finding, along with inhibition of Dpm1 rescuing disease phenotypes, suggest that the more likely mechanism may be increased substrate availability for Alg7.

*RPS9B* is not a known physical or genetic interactor of *SDO1*. *TIF6*, on the other hand, is a known genetic interactor of *SDO1*, and *RIA1* is a known genetic and physical interactor (Costanzo et al., 2016; Krogan et al., 2006). The protein products of these genes work together closely; Sdo1 and Ria1 function together to promote the release of Tif6 from the 60S ribosomal subunit so it can associate with the 40S subunit, which contains Rps9b (Finch et al., 2011; Menne et al., 2007). The physical interaction between *SDO1* and *RIA1* could account for the allele-specificity we observed in suppression. *RIA1* mutations occurred only in *sdo1*-E44G strains during the evolution experiment, and we found that these evolved mutations only increased fitness in this background (Supplemental Fig. 1). E44G is predicted to affect surface electrostatic potential or alter surface epitopes of the protein product, suggesting that it is directly involved in Ria1 associations (Shammas et al., 2005). Mutations occurring in *RIA1* could therefore be compensating by stabilizing this protein-protein interaction. In contrast, *sdo1*-K62* produces a truncated protein that cannot make the same contacts, so *RIA1* mutations have little to no effect.

*RPS9B* mutations also demonstrated allele-specificity of suppression, with mutations evolving in *sdo1*-K62* backgrounds, but not in *sdo1*-E44G. From the competitive fitness assay, we found that both evolved *RPS9B* mutations caused fitness defects in wildtype backgrounds, with defects becoming even more severe in combination with *sdo1*-E44G. Mutations in *RPS9B* could be altering association dynamics in ribosomal subunit assembly, decreasing the requirement of *SDO1* for *TIF6* release. Finally, *TIF6* rescued defects from both *SDO1* disease-associated alleles, in line with previous findings from patient data and human cell lines in which reduced expression of the human ortholog, *EIF6*, or mutations disrupting Eif6 binding enabled ribosomal subunit assembly and resulted in more mild disease symptoms (Kennedy et al., 2021; Tan et al., 2021; Valli et al., 2019). In this case, the specific allele of *SDO1* has less impact on suppression, as the reduced binding of Tif6 circumvents the need for a functional Sdo1 to release Tif6 from the 60S subunit.

Identifying genetic suppressors has the potential to inform strategies for mitigating disease and for drug repurposing (Jonker et al., 2024; Roessler et al., 2021). For example, experimental evolution of yeast strains harboring variants associated with Wiskott-Aldrich syndrome found that loss-of-function mutations in calcineurin compensated for disease phenotypes, suggesting cyclosporin A as a potential treatment (Filteau et al., 2015). Here, we demonstrate the strength of yeast experimental evolution coupled with whole genome sequencing to identify allele-specific suppressors in yeast models of rare human disease. We expanded this framework (Vignogna et al., 2022) to an additional 11 disease alleles across six genes, identifying both known and unknown suppressors of *ALG7* and *SDO1*.

There are more than 7,000 rare diseases, with over 4,000 known causative genes (Kachroo et al., 2022; Lee et al., 2020; Stoller, 2018). Individual rare diseases, by definition, affect less than 200,000 individuals, but collectively rare diseases affect at least 300 million people worldwide (Nguengang Wakap et al., 2020). Despite this, 95% of these conditions still have no available treatments (Nguengang Wakap et al., 2020; Paltenghi-Peceva et al., 2026). Given that 20-25% of human genes have conserved yeast orthologs, we estimate 800-1,000 rare disease-associated genes to which we could apply this approach (Lee et al., 2020). Ultimately, experimental evolution has the potential to identify allele-specific genetic suppressors of a large number of rare human diseases.

## METHODS

### Yeast strain constructions

Haploid and diploid strains used in this study are S288C derivatives with *ura3Δ*0 auxotrophy. All cultures were grown in nutrient-rich media (YPD; 1% yeast extract, 2% peptone, 2% dextrose) at 30°C unless otherwise specified.

To construct strains harboring human disease-associated alleles, we utilized a two-plasmid CRISPR-Cas9 editing system. We PCR amplified 500 bp, commercially synthesized dsDNA fragments (Integrated DNA Technologies, Coralville, IA, USA) containing both a 20 bp guide sequence and approximately 300 bp repair template with the mutation of interest. These gBlocks also encode AscI and NotI-HF restriction sites for ligation into a plasmid, as well as sgRNA scaffolding and a transcription terminator.

Amplified and restriction-digested gBlocks were ligated into a 2-micron plasmid derived from pRS425, also containing *LEU2* and KanMX markers and LexA sites. The second plasmid is a CEN/ARS plasmid derived from pKR193 (Roy et al., 2018), encoding a galactose-inducible Cas9 and a LexA-Fkh1p fusion protein, plus KlURA3 and HphMX markers. We co-transformed these two plasmids into both *MAT***a** and *MAT*α haploid strains and induced Cas9 expression by plating the transformants onto YPGal (1% yeast extract, 2% peptone, 2% galactose). After Cas9 cuts the target DNA based on the guide sequence, Fkh1p within the LexA-Fkh1p fusion protein binds to the double-stranded DNA break. Simultaneously, the LexA portion of the fusion protein binds to the LexA sites within the 2-micron plasmid. Through these interactions, the repair template is brought into close proximity of the cut site, promoting its use in homology-directed repair of the double-stranded break and thus introducing the human disease-allele changes into the yeast ortholog (Roy et al., 2018). We confirmed successful gene editing by Sanger sequencing (Psomagen, Rockville, MD, USA).

We mated each resulting strain with the parental strains, now containing a plasmid with a NatMX marker, and selected for diploids based on resistance to both Hygromycin (300 µg/ml) and CloNat (100 µg/ml). This allowed us to generate heterozygous diploids, as well as perform tetrad dissections to obtain any haploids in which only one mating type was successfully edited previously. Haploids with confirmed mutations were mated to generate homozygous diploids and compound heterozygotes when applicable. We then selected for plasmid loss by streaking them to 5-FOA (0.67% yeast nitrogen base, 0.1% 5-fluoroortic acid, 0.002% uracil, 2% dextrose) and again confirmed mutations in diploids by Sanger sequencing.

To reconstruct strains harboring both the original disease-associated alleles and evolved mutations identified by whole-genome sequencing analysis, we used the same two-plasmid system with gBlocks designed to generate the evolved mutations and co-transformed into the original *MAT*α background strain. After successful editing was confirmed by Sanger sequencing, we mated these strains with the ancestral *MAT***a** disease strains, now with a NatMX marker. Again, we selected for diploids with resistance to Hygromycin and CloNat, then selected for spontaneous plasmid loss on 5-FOA.

### Tetrad dissections

Heterozygous diploid strains were sporulated in SPO++ (1.5% potassium acetate, 0.25% yeast extract, 0.25% dextrose, 0.002% histidine, 0.002% tryptophan) on a roller-drum at room temperature for approximately five days. For each strain, 10 to 20 tetrads were dissected on YPD agar and incubated at 30°C for 48 hours.

For reconstructions of ancestral disease-associated alleles with suppressor alleles, we assessed the segregation of large and small colony sizes among four-spore tetrads. Based on the segregation patterns, we calculated the likelihood of the data under models of suppression (P(data | sup)) and no suppression (P(data | ∼sup)). We then calculated log_10 l_ikelihood ratio (LLR) as log_10[_P(data | sup) / P(data | ∼sup)], where positive values favor suppression and negative values favor no suppression. Exact multinomial p-values were calculated in R to assess statistical significance.

### Quantifying growth rate

From each tetrad dissection plate, we chose four 4-spore viable tetrads, inoculating each well of 96-well plates with an individual spore in 150 µl of YPD. 96-well plates were incubated for 24 hours at 30°C. The cultures were then diluted 1:1,000 and we used a Tecan Infinite M Plex plate reader to measure optical density (OD_600)_ over the course of 24 hours at 30°C. Reads were taken every 15 minutes with 4 mm orbital shaking for two minutes prior, and with multiple reads per well (3 x 3 grid). Plots of growth curves were generated in R.

### Experimental evolution

Single colonies from each strain were used to inoculate 3 ml of YPD and these cultures were grown to saturation on a roller-drum at 30°C before being distributed into four 96-well plates. Heterozygous diploids were distributed in replicates of three, while haploids, homozygous diploids, and compound heterozygous diploids were done in replicates of eight. Each plate was incubated at 30°C without shaking for 24 hours. We used a BiomekFX liquid handler to dilute each population 1:32 every 24 hours by transferring 4 µl of culture into 125 µl of YPD plus 100 µg/ml ampicillin and 25 µg/ml tetracycline for prevention of bacterial contamination. This dilution factor allows for approximately five generations of growth per day and we proceeded for 1,000 generations. While this experimental approach has previously been done with a 1:1024 dilution for 10 generations of growth per day (Fisher et al., 2019; Martínez et al., 2023; Vignogna et al., 2022), the *sdo1*-K62* strain grew too slowly to be maintained with this dilution. We therefore adjusted to a 1:32 dilution in order to maintain the slowest growing populations. Once a week, we added 50 µl of 60% glycerol to each well and froze the plates at -80°C.

### Competitive fitness assays

We measured fitness effects at generations 0, 300, 700, and 1,000 of the evolution experiment using a flow-cytometry based competitive fitness assay, as previously described (Buskirk et al., 2017). Rather than the usual 1:1 mix used for this assay, the experimental strains were mixed 75:25 with a fluorescently-labeled (ymCitrine) version of the wildtype background strain. This ratio adjustment was based on previous work where an exceedingly slow-growing disease strain, much like our *sdo1*-K62* strain, was immediately outcompeted by the reference, preventing accurate measurements of fitness effects (Vignogna et al., 2022). Cocultures were propagated under the same conditions as the evolution experiment for 30 generations, with 4 µl transferred into 70 µl PBS each day to be run on the BD FACSCanto II flow cytometer. A minimum of 30,000 events were recorded per sample. Biological replicates were grouped for analysis at generation 0, with mean and standard deviation reported, but were treated as individual populations for subsequent generations as they evolved independently. Selection coefficients were estimated as the slope of ln(p_ref/p_population) versus generations. Analyses were performed in Flowjo and R, with plots generated using ggplot2.

For reconstructions of ancestral disease-associated alleles with suppressor alleles, we calculated the mean fitness effects of spores with the disease-associated allele alone, those with both the disease and suppressor allele (double mutants), and spores with wildtype alleles. We could not always distinguish between wildtype spores with and without the suppressor, as they often had comparable fitness values. In these cases, statistically half of the spores are true wildtype and half are wildtype plus the suppressor. Using mean fitness effects (F), we calculated percent restoration as (F_Double Mutant –_ F_Disease Allele)_/(F_Wildtype –_ F_Disease Allle)_*100.

### Whole-genome sequencing

We isolated a single clone from each evolved population at generation 1,000 and extracted genomic DNA by enzymatic lysis with zymolyase, followed by ethanol precipitation and RNase A treatment. Nextera libraries were prepared as described previously (Buskirk et al., 2017). Pooled libraries were sequenced by the Lewis-Sigler Institute for Integrative Genomics at Princeton University using an Illumina NovaSeq 6000 sequencer. Haploids and diploids were sequenced in two separate batches.

### Sequencing analysis

For analysis of whole-genome sequencing data, we followed a previously described pipeline (Martínez & Lang, 2023) with slight adjustments. We trimmed Illumina 150 bp paired-end reads with Trimmomatic (v0.39) and used BWA (v0.7.17) to align the reads with the *S. cerevisiae* S288C reference genome (downloaded February 2025 from the Yeast PacBio resource: https://yjx1217.github.io/Yeast_PacBio_2016/data/). Samtools (v1.19.2) removed PCR duplicates and calculated per-site coverage. Variants were called with GATK (v4.5.0.0), filtered by VCFtools (v0.1.16) to remove low-quality and common calls, and annotated with OpenJDK (v17.0.11_9) and SnpEff (v5.0). Calls were confirmed manually by viewing BAM files in IGV. We calculated the probability of observing a given number of nonsynonymous mutations in each gene using a Poisson distribution.

For genes with a significant number of nonsynonymous mutations, we assessed whether these mutations were more consistent with selection for loss-of-function (LOF) or non-loss-of-function (non-LOF) by performing a Bayesian model comparison. We calculated the likelihood of the observed frequencies of missense versus nonsense and frameshift mutations under LOF and non-LOF hypotheses, then used these likelihoods to calculate posterior probabilities of the two hypotheses (Martínez & Lang, 2023). We calculated log_10 l_ikelihood ratios as log_10[_LOF/non-LOF].

### Quantifying alpha-factor secretion by halo assay

We used a mating type halo assay to examine the release of alpha factor in *ALG7* mutant strains. We mixed 90 µl of YPD with 10 µl of a saturated culture of *MAT***a** mating type tester, which has a mutation in *SST2* leading to elevated sensitivity to mating pheromone. All 100 µl were spread on YPD agar using glass beads and allowed to dry for approximately ten minutes before pinning strains of interest onto the plates. Plates were incubated for 48 hours at 30°C and photos were taken using an Epson scanner. We used Fiji to measure the diameter of halos which formed around *MAT*α cells, as well as diameter of the colonies themselves. Each diameter was measured three times, and the average was used to determine the area of the circle. Final area was calculated by subtracting the area of the colony from the area of the halo.

## ACKNOWLEDGEMENTS

We thank members of the Lang Lab for comments on the manuscript.

## DATA AVAILABILITY

The short-read sequencing data reported in this study have been deposited to the NCBI BioProject database, accession number PRJNA1566777.

## AUTHOR CONTRIBUTIONS

Conceptualization: GIL

Experiments: BD

Analysis: BD

Writing: BD, GIL

Editing: BD, GIL

Funding Acquisition: GIL

