## Supplementary Figure 1 for "Evolutionary rescue of rare human genetic disorders in yeast models"

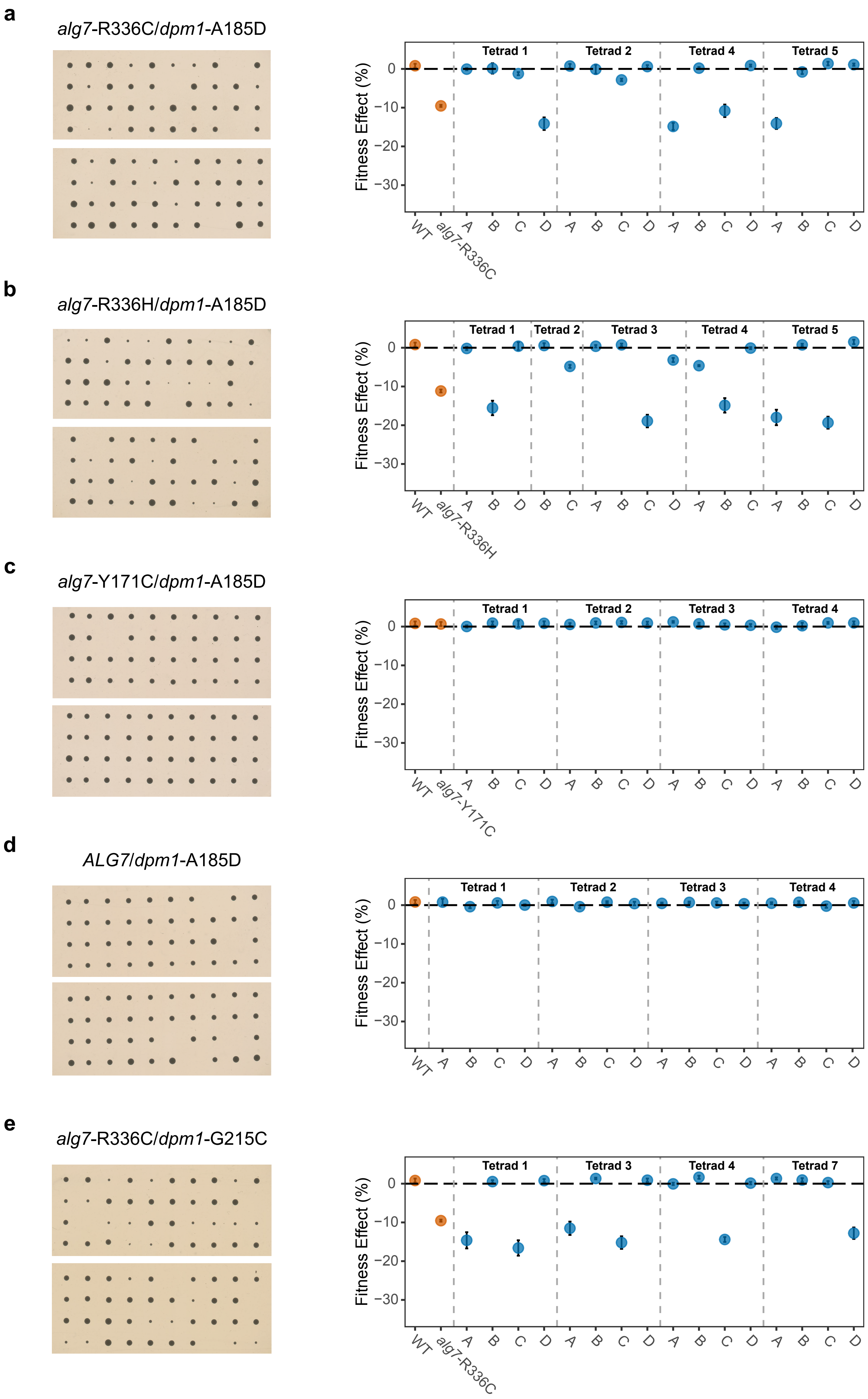
a
alg7-R336C/dpm1-A185D
Fitness Effect (%)
Tetrad 1
Tetrad 2
Tetrad 4
Tetrad 5
0
−10
−20
−30
WT
alg7-R336C
A B C D A B C D A B C D A B C D
b
alg7-R336H/dpm1-A185D
Fitness Effect (%)
Tetrad 1
Tetrad 2
Tetrad 3
Tetrad 4
Tetrad 5
0
−10
−20
−30
WT
alg7-R336H
A B D B C A B C D A B C A B C D
c
alg7-Y171C/dpm1-A185D
Fitness Effect (%)
Tetrad 1
Tetrad 2
Tetrad 3
Tetrad 4
0
−10
−20
−30
WT
alg7-Y171C
A B C D A B C D A B C D A B C D
d
ALG7/dpm1-A185D
Fitness Effect (%)
Tetrad 1
Tetrad 2
Tetrad 3
Tetrad 4
0
−10
−20
−30
WT
A B C D A B C D A B C D A B C D
e
alg7-R336C/dpm1-G215C
Fitness Effect (%)
Tetrad 1
Tetrad 3
Tetrad 4
Tetrad 7
0
−10
−20
−30
WT
alg7-R336C
A B C D A B C D A B C D A B C D

**f**

*alg7*-R336H/*dpm1*-G215C

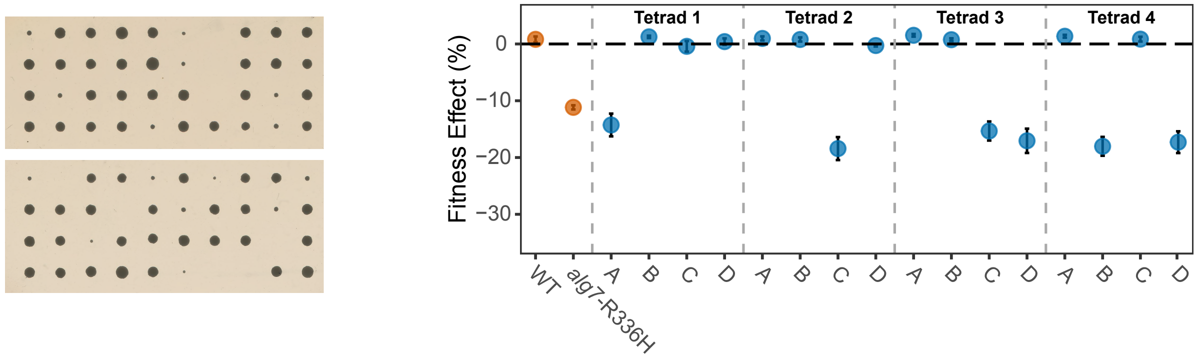


**g**

*alg7*-Y171C/*dpm1*-G215C

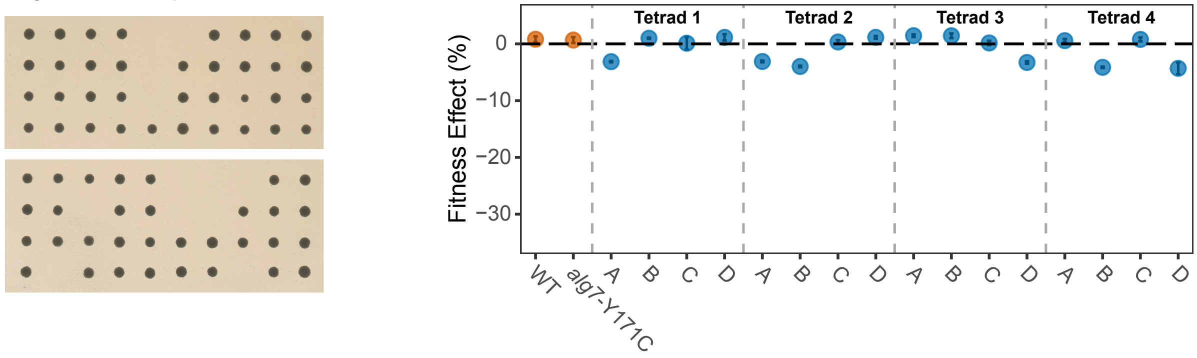


**h**

*ALG7*/*dpm1*-G215C

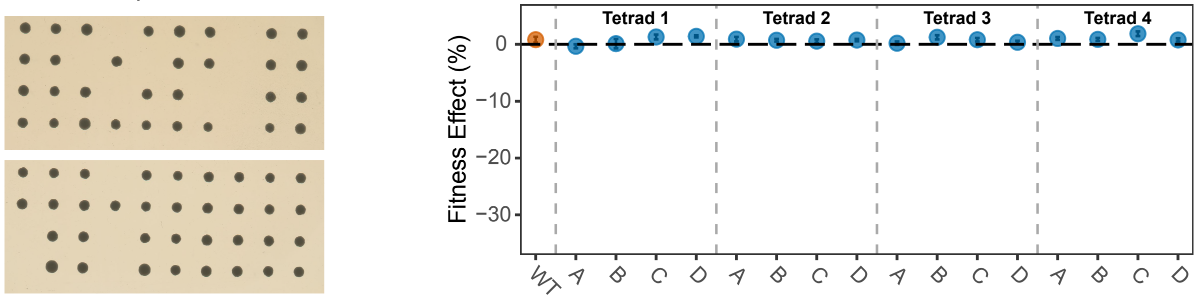


**i**

*alg7*-R336C/*psa1*-D48Y

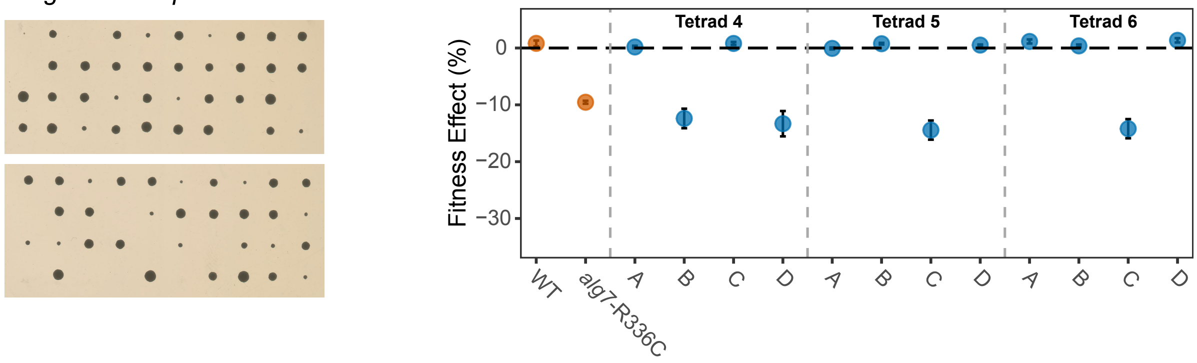


**j**

*alg7*-R336H/*psa1*-D48Y

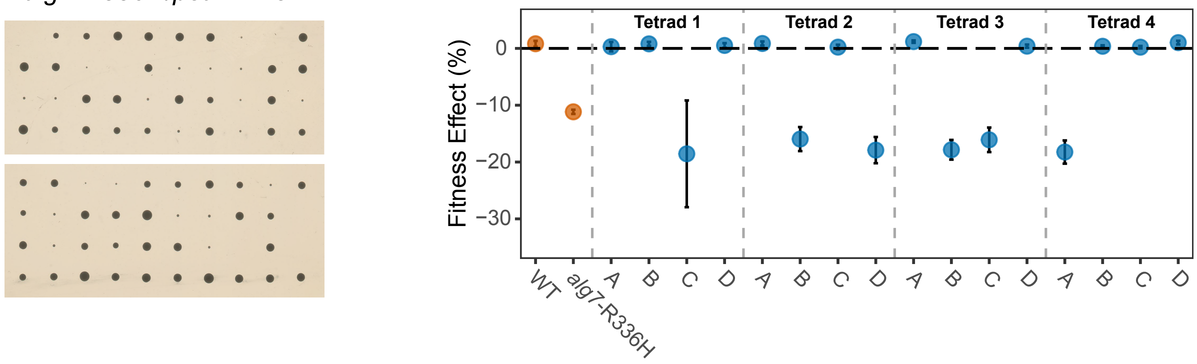

**k**

*alg7*-Y171C/*psa1*-D48Y

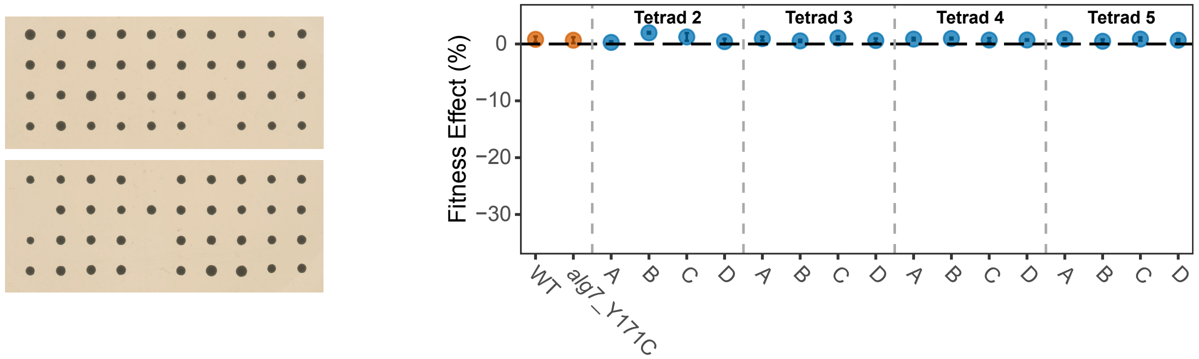


**l**

*ALG7*/*psa1*-D48Y

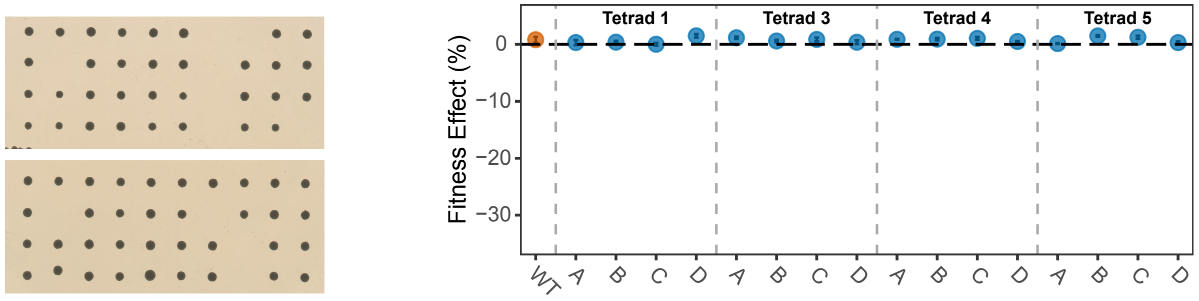


**m**

*alg7*-R336C/*psa1*-W311R

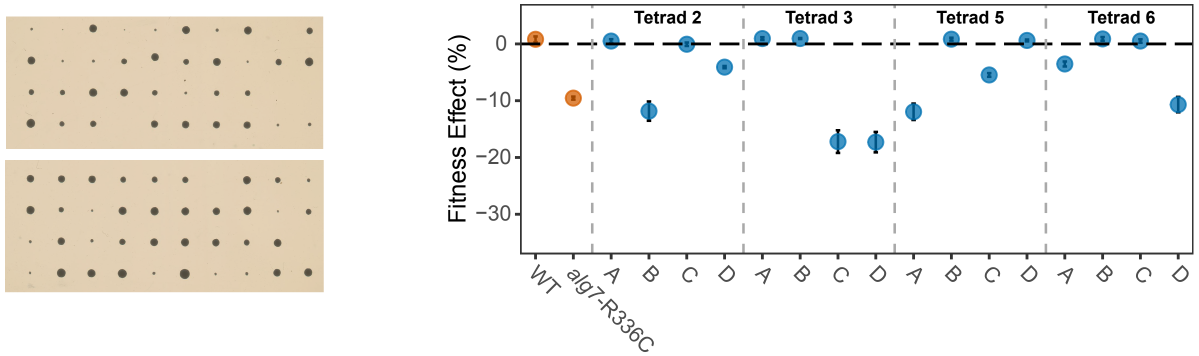


**n**

*alg7*-R336H/*psa1*-W311R

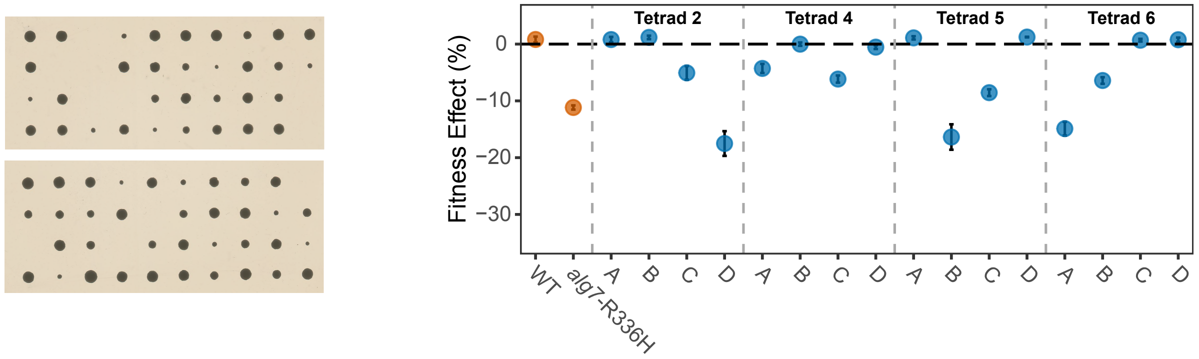


**o**

*alg7*-Y171C/*psa1*-W311R

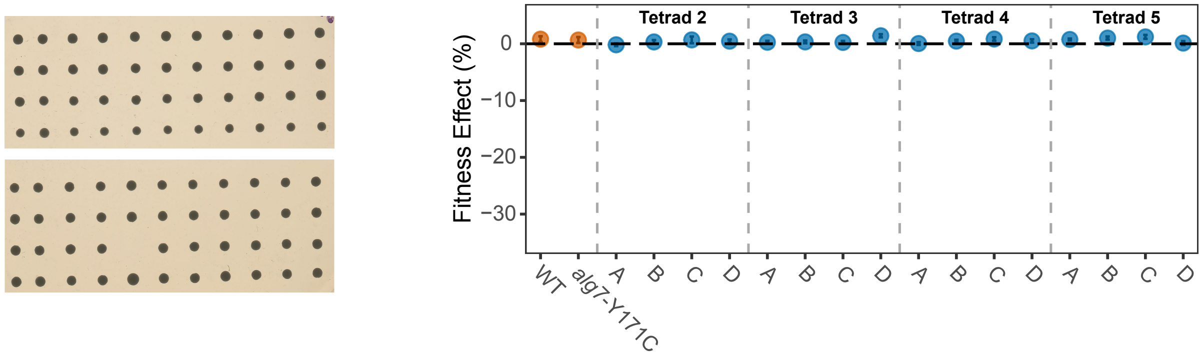

**p**

*ALG7*/*psa1*-W311R

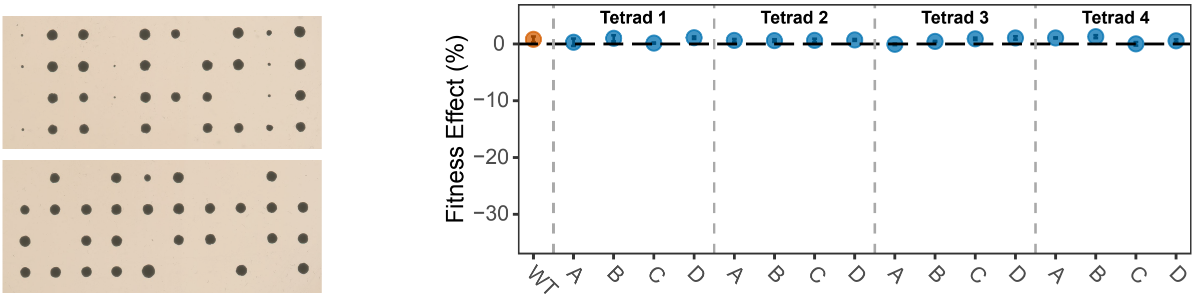


**q**

*alg7*-R336C/*nnk1*-R326K

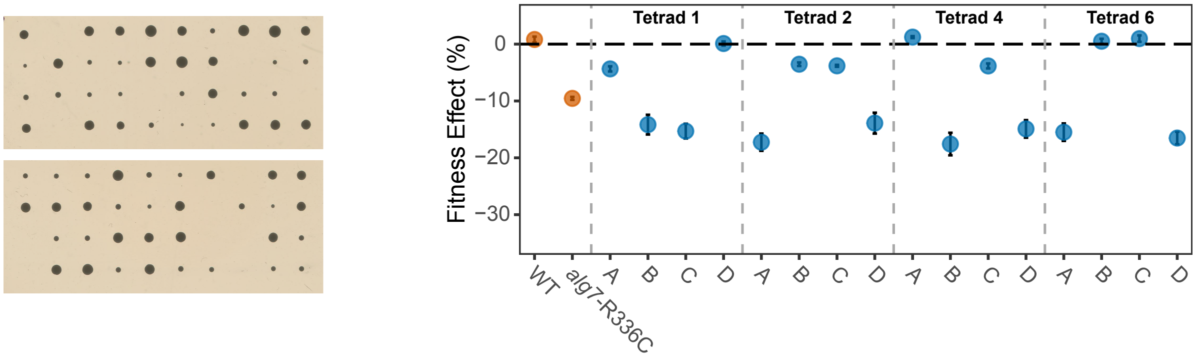


**r**

*alg7*-R336H/*nnk1*-R326K

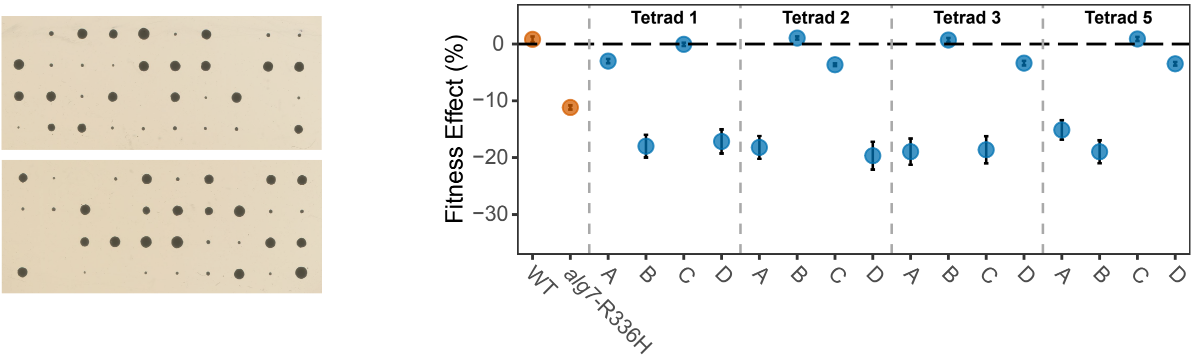


**s**

*alg7*-Y171C/*nnk1*-R326K

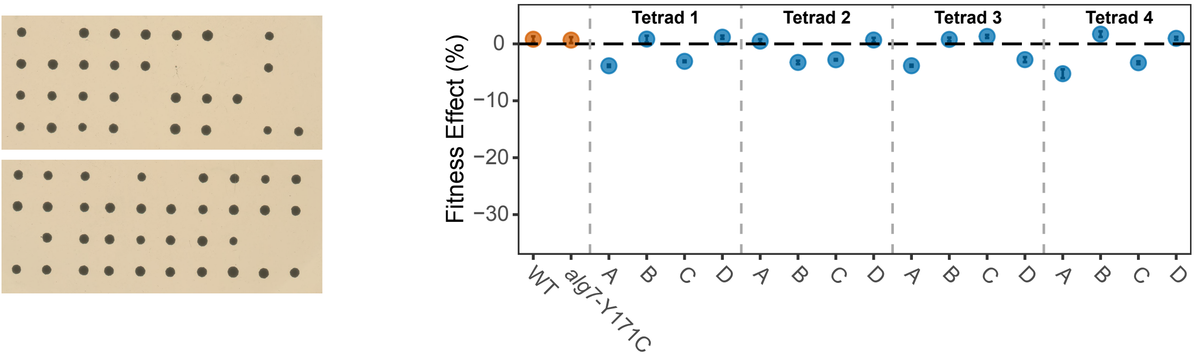


**t**

*ALG7*/*nnk1*-R326K

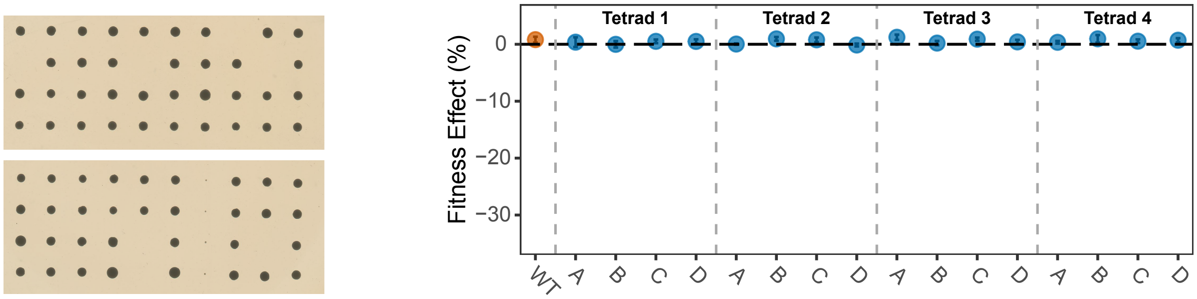

u

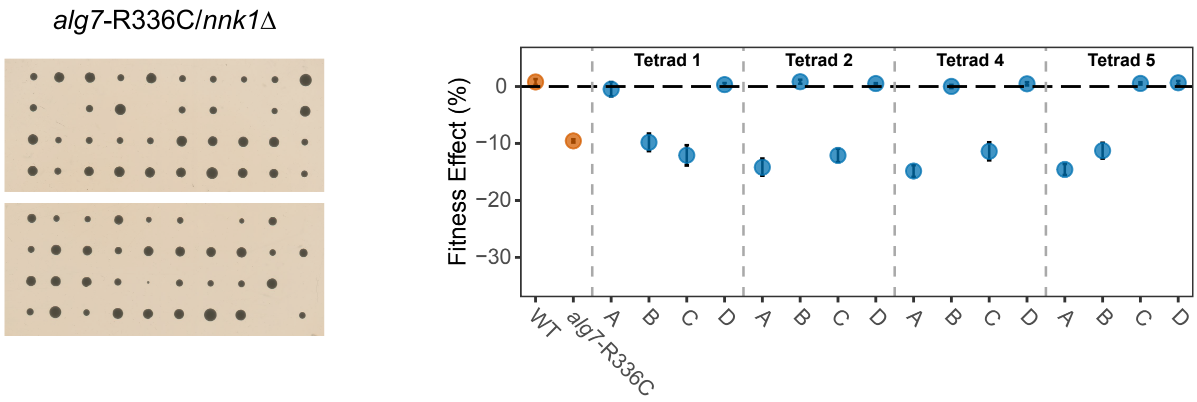

*alg7*-R336C/*nnk1*Δ
Tetrad 1
Tetrad 2
Tetrad 4
Tetrad 5
Fitness Effect (%)
0
−10
−20
−30
WT
alg7-R336C
A B C D A B C D A B C D A B C D


v

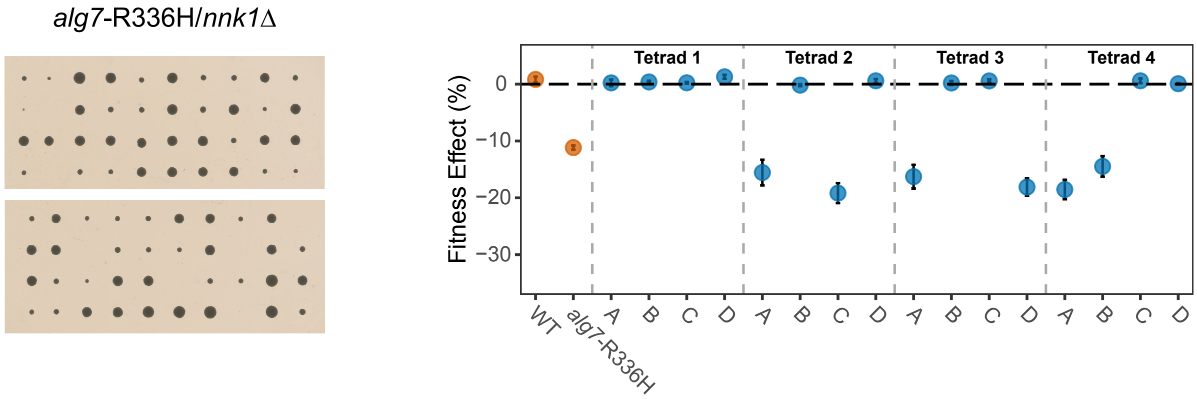

*alg7*-R336H/*nnk1*Δ
Tetrad 1
Tetrad 2
Tetrad 3
Tetrad 4
Fitness Effect (%)
0
−10
−20
−30
WT
alg7-R336H
A B C D A B C D A B C D A B C D


w

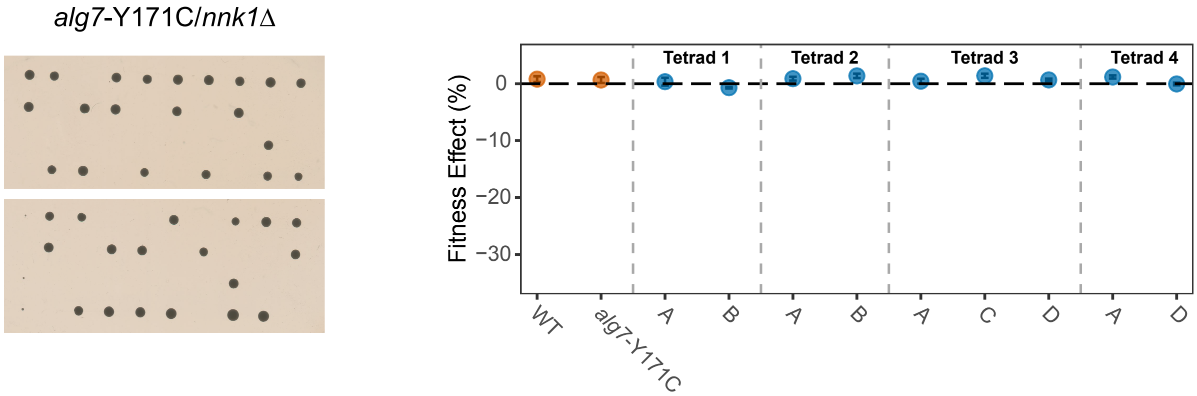

*alg7*-Y171C/*nnk1*Δ
Tetrad 1
Tetrad 2
Tetrad 3
Tetrad 4
Fitness Effect (%)
0
−10
−20
−30
WT
alg7-Y171C
A B A B A C D A D


x

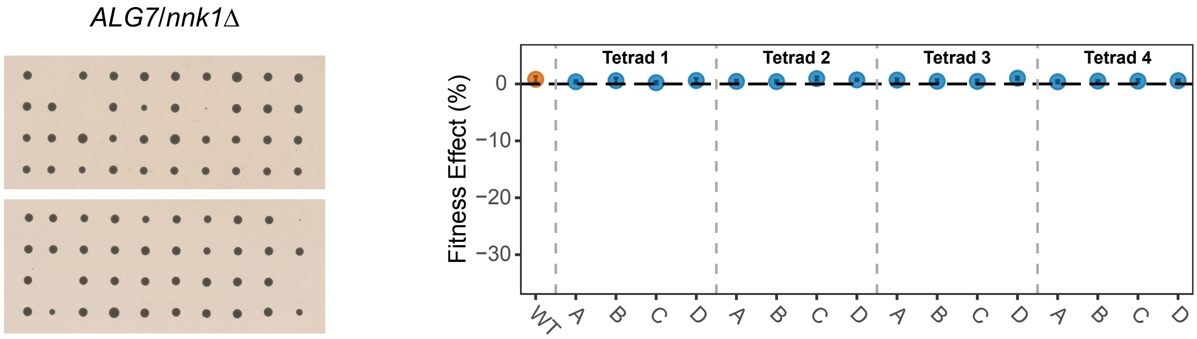

*ALG7*/*nnk1*Δ
Tetrad 1
Tetrad 2
Tetrad 3
Tetrad 4
Fitness Effect (%)
0
−10
−20
−30
WT
A B C D A B C D A B C D A B C D


y

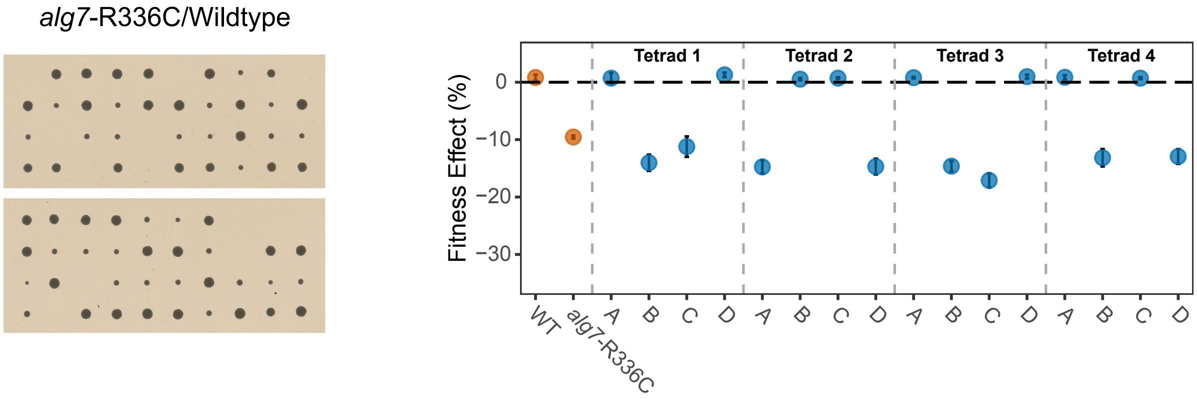

*alg7*-R336C/Wildtype
Tetrad 1
Tetrad 2
Tetrad 3
Tetrad 4
Fitness Effect (%)
0
−10
−20
−30
WT
alg7-R336C
A B C D A B C D A B C D A B C D

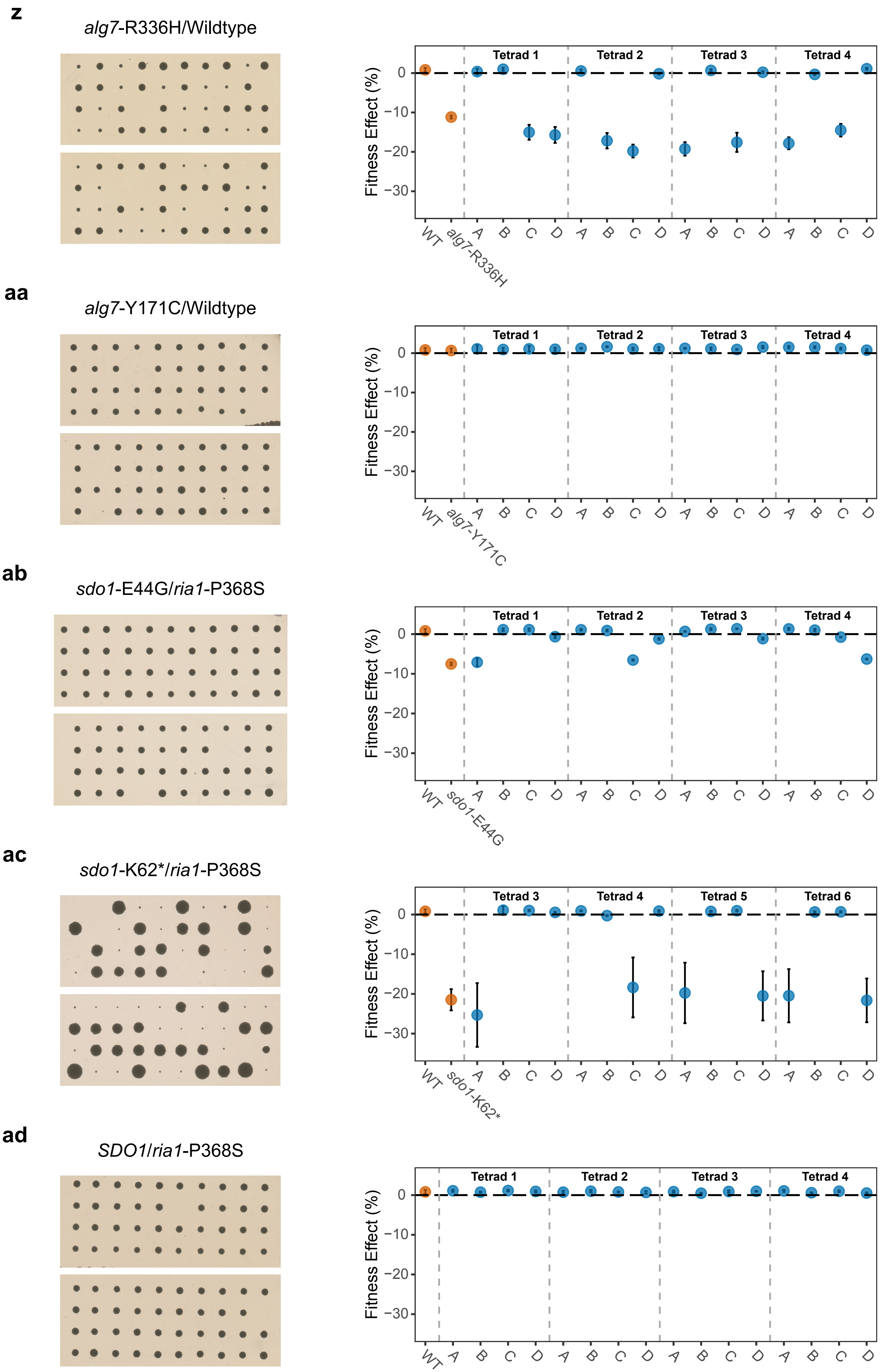
z
alg7-R336H/Wildtype
Tetrad 1
Tetrad 2
Tetrad 3
Tetrad 4
Fitness Effect (%)
0
−10
−20
−30
WT
alg7-R336H
A B C D A B C D A B C D A B C D
aa
alg7-Y171C/Wildtype
Tetrad 1
Tetrad 2
Tetrad 3
Tetrad 4
Fitness Effect (%)
0
−10
−20
−30
WT
alg7-Y171C
A B C D A B C D A B C D A B C D
ab
sdo1-E44G/ria1-P368S
Tetrad 1
Tetrad 2
Tetrad 3
Tetrad 4
Fitness Effect (%)
0
−10
−20
−30
WT
sdo1-E44G
A B C D A B C D A B C D A B C D
ac
sdo1-K62*/ria1-P368S
Tetrad 3
Tetrad 4
Tetrad 5
Tetrad 6
Fitness Effect (%)
0
−10
−20
−30
WT
sdo1-K62*
A B C D A B C D A B C D A B C D
ad
SDO1/ria1-P368S
Tetrad 1
Tetrad 2
Tetrad 3
Tetrad 4
Fitness Effect (%)
0
−10
−20
−30
WT
A B C D A B C D A B C D A B C D

**ae**

*sdo1*-E44G/*ria1*-Y991C

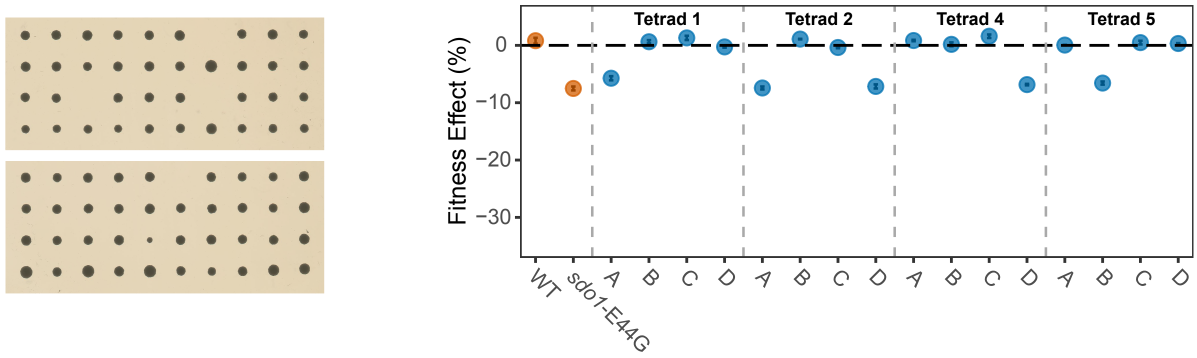


**af**

*sdo1*-K62*/*ria1*-Y991C

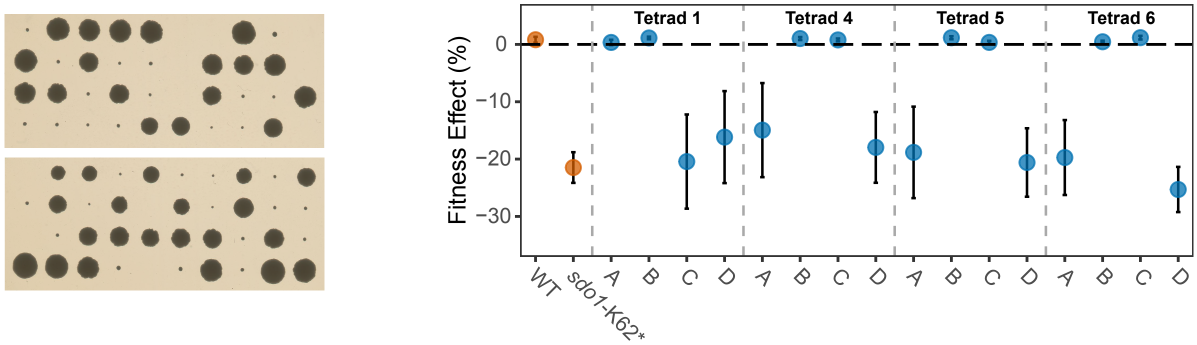


**ag**

*SDO1*/*ria1*-Y991C

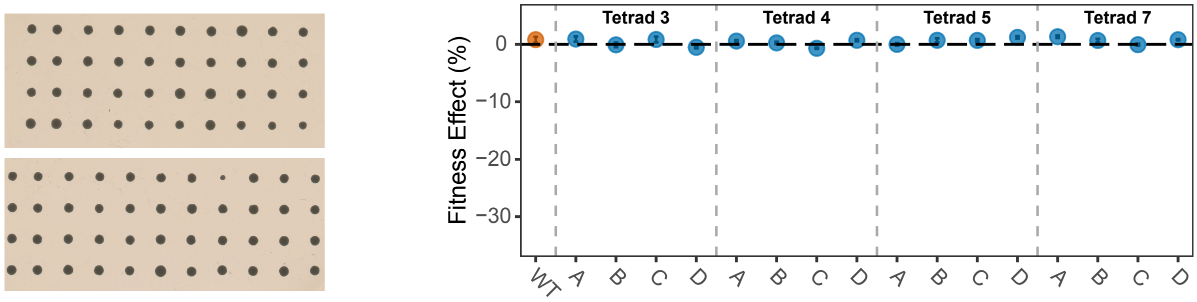


**ah**

*sdo1*-E44G/*rps9b*-K92Δ

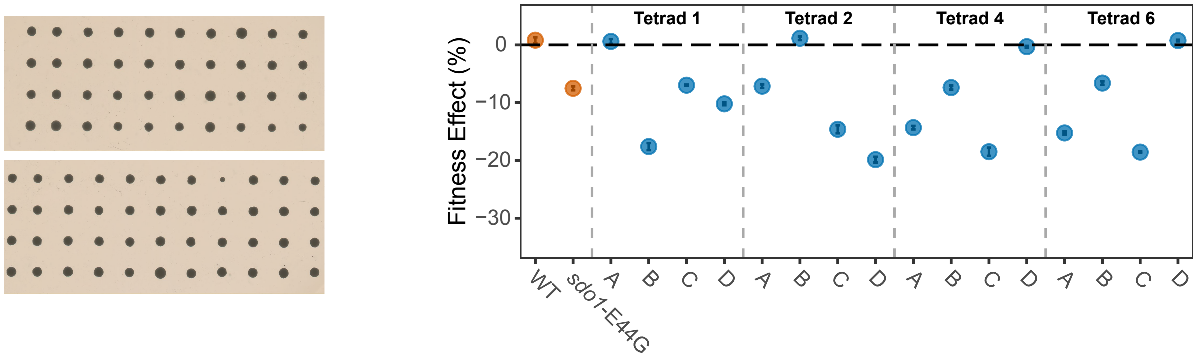


**ai**

*sdo1*-K62*/*rps9b*-K92Δ

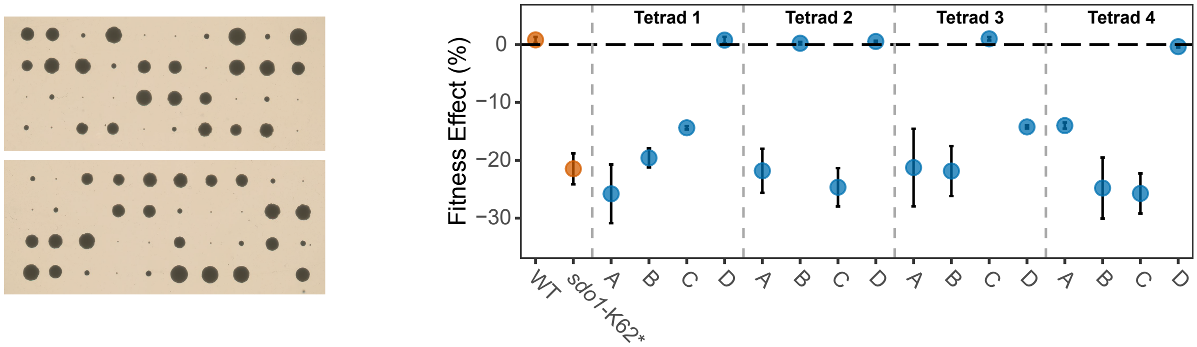

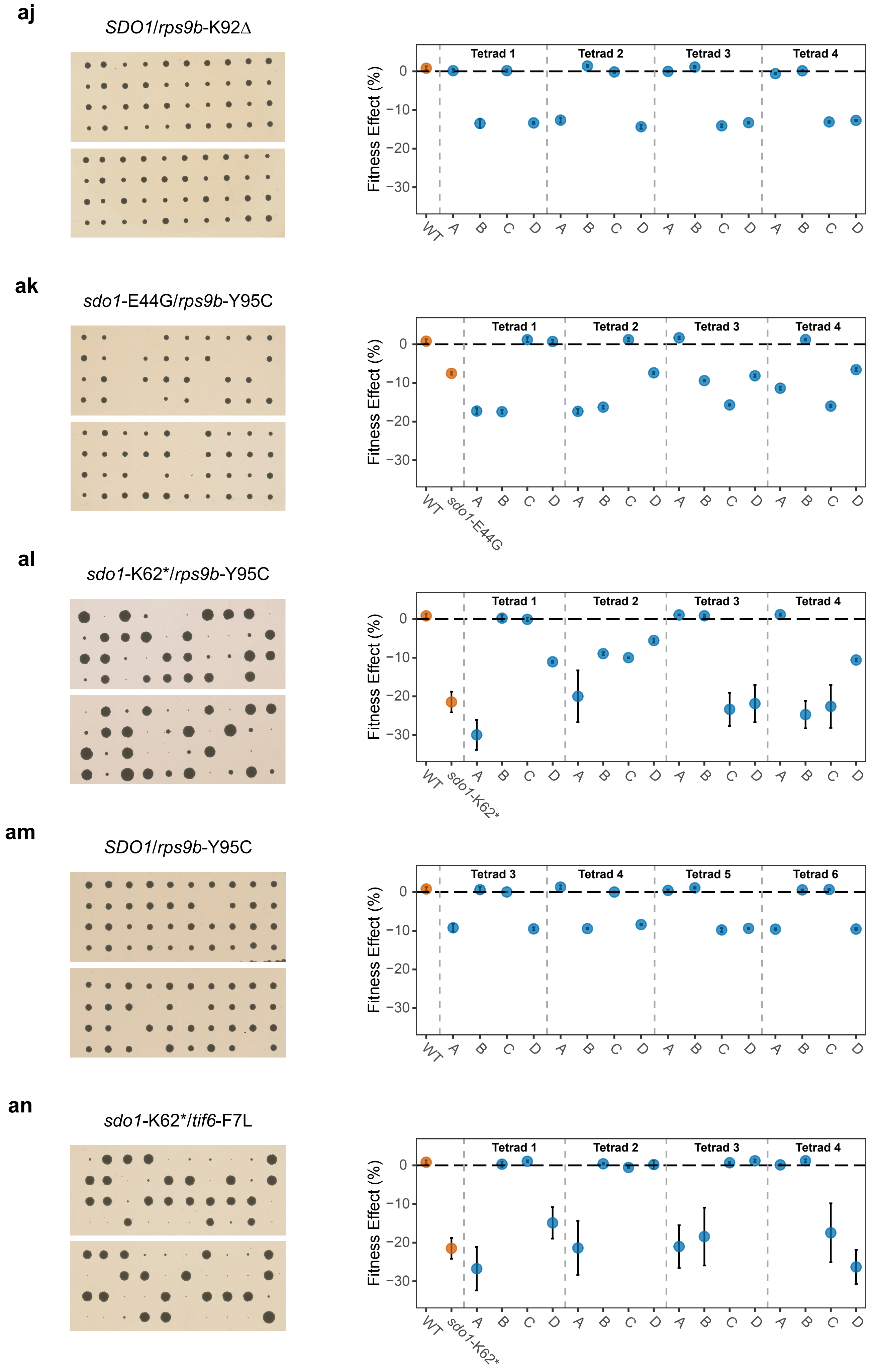
aj
SDO1/rps9b-K92Δ
Fitness Effect (%)
Tetrad 1
Tetrad 2
Tetrad 3
Tetrad 4
0
−10
−20
−30
WT A B C D A B C D A B C D A B C D
ak
sdo1-E44G/rps9b-Y95C
Fitness Effect (%)
Tetrad 1
Tetrad 2
Tetrad 3
Tetrad 4
0
−10
−20
−30
WT sdo1-E44G A B C D A B C D A B C D A B C D
al
sdo1-K62*/rps9b-Y95C
Fitness Effect (%)
Tetrad 1
Tetrad 2
Tetrad 3
Tetrad 4
0
−10
−20
−30
WT sdo1-K62* A B C D A B C D A B C D A B C D
am
SDO1/rps9b-Y95C
Fitness Effect (%)
Tetrad 3
Tetrad 4
Tetrad 5
Tetrad 6
0
−10
−20
−30
WT A B C D A B C D A B C D A B C D
an
sdo1-K62*/tif6-F7L
Fitness Effect (%)
Tetrad 1
Tetrad 2
Tetrad 3
Tetrad 4
0
−10
−20
−30
WT sdo1-K62* A B C D A B C D A B C D A B C D

**ao**

*SDO1*/*tif6*-F7L

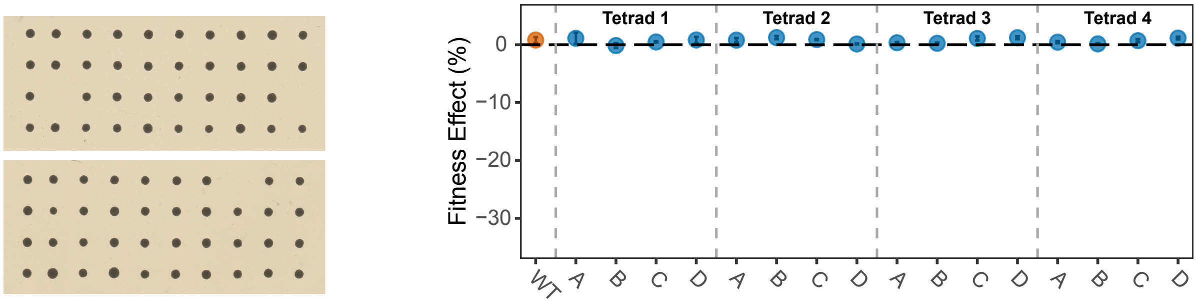


**ap**

*sdo1*-E44G/*tif6*-I58T

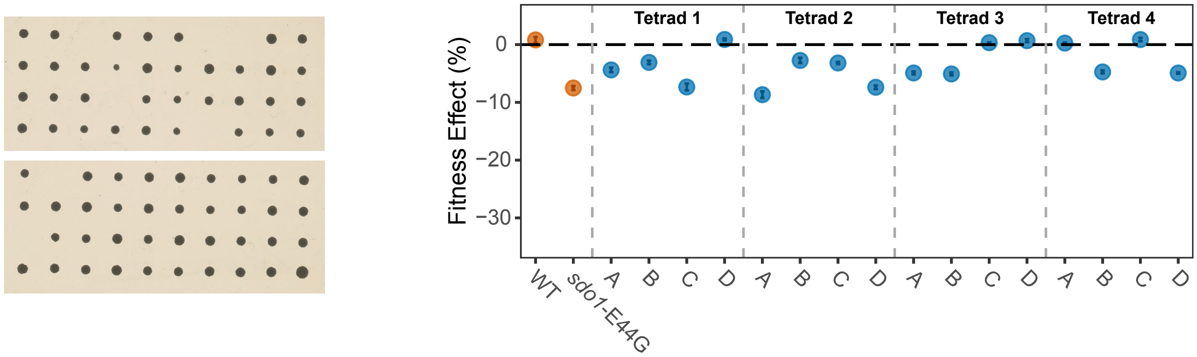


**aq**

*sdo1*-K62*/*tif6*-I58T

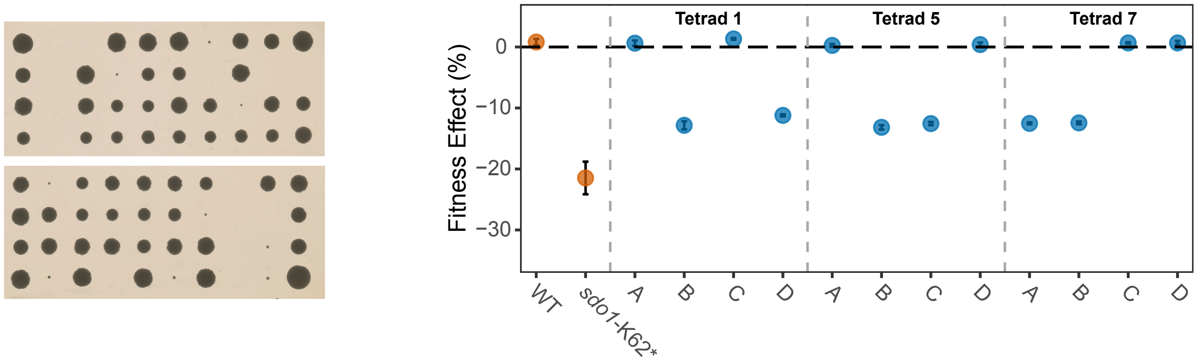


**ar**

*SDO1*/*tif6*-I58T

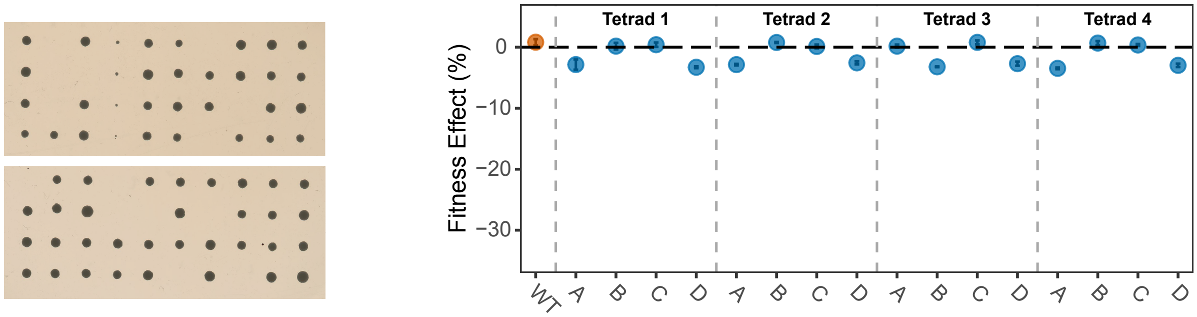


**as**

*sdo1*-E44G/Wildtype

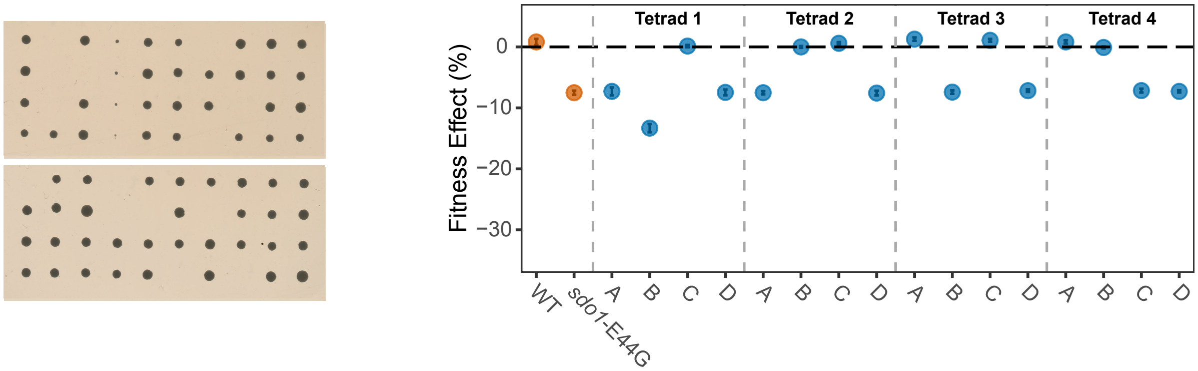

**at**

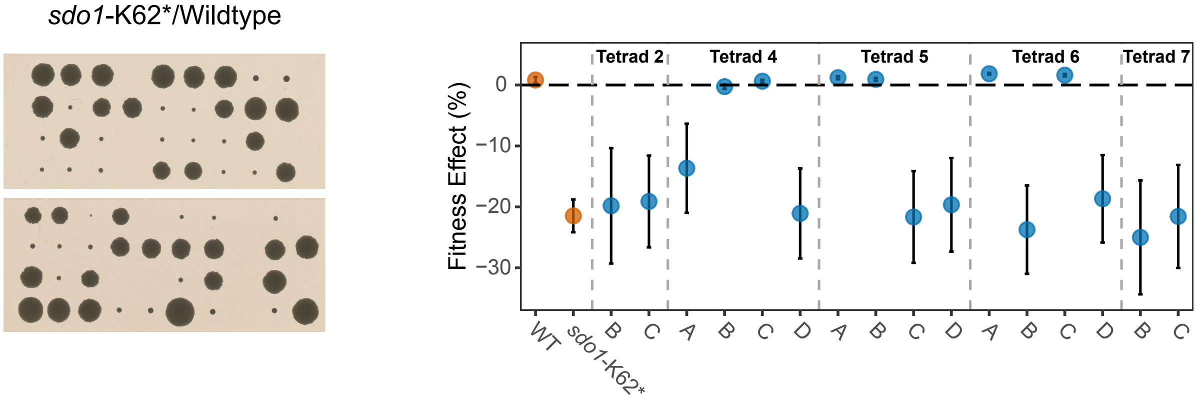


**au**

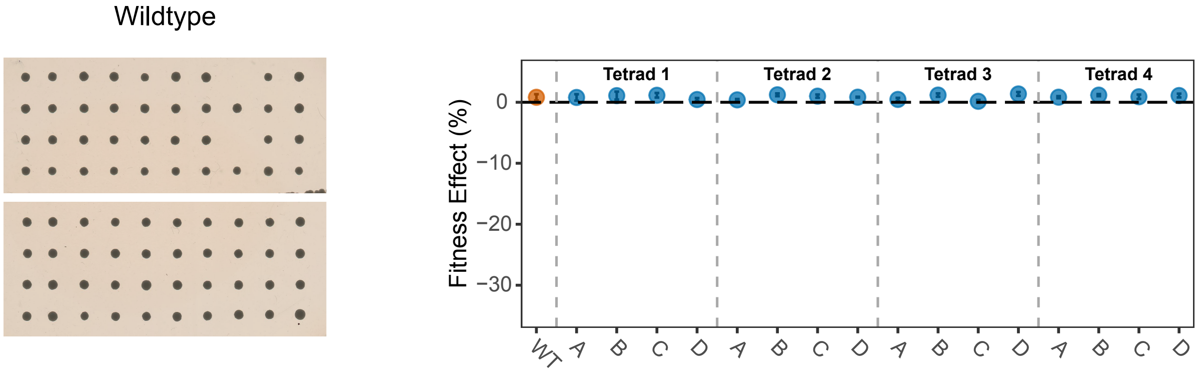


**Supplemental Fig. 1.** Tetrad dissections and fitness effects for heterozygous reconstructions containing ancestral disease-associated alleles and putative suppressors. Plate pictures for tetrad dissections were taken after two days of growth, with the exception of those possessing *sdo1*-K62* spores, which required three to six days to become visible. Fitness effects were measured by competitive fitness assay. We show four spores from one tetrad per cross, with error bars representing the standard error of slope. We include a comparison of wildtype and averaged *MAT***a** and *MAT*α haploids from Generation 0 of the evolution experiment, with error bars indicating the standard deviation of the mean (Supplemental Dataset 1).  Reconstructuon fitness data and log-likelihood calculations for suppression based on colony size segregation patterns can be found in Supplemental Dataset 3.
